# Evaluating drivers of spatiotemporal individual condition of a bottom-associated marine fish

**DOI:** 10.1101/2022.04.19.488709

**Authors:** Max Lindmark, Sean C. Anderson, Mayya Gogina, Michele Casini

**Author notes:** Author to whom correspondence should be addressed. Max Lindmark, Swedish University of Agricultural Sciences, Department of Aquatic Resources, Institute of Marine Research, Turistgatan 5, 453 30 Lysekil, Sweden.

## Abstract

An organism’s body condition describes its mass given its length and is often positively associated with fitness. The condition of Atlantic cod (*Gadus morhua*) in the Baltic Sea has declined dramatically since the early 1990s, possibly due to increased competition, food limitation, and hypoxia. However, the effect of biotic and abiotic variables on body condition have not been evaluated at local scales, which is important given spatial heterogeneity. We evaluate changes in distribution, experienced environmental conditions, and individual-level condition of cod in relation to covariates at different spatial scales using geostatistical models with spatial and spatiotemporal random effects. Oxygen, sprat biomass, and temperature were positively related to condition, and depth negatively associated, but the effect sizes of these variables were small—spatial and spatiotemporal latent variables explained almost five times more variation than fixed effects. We also show that accounting for the heterogenous distribution of cod leads to both lover levels and steeper trends over time in experienced oxygen compared to those in the environment. Understanding the drivers of spatiotemporal variation in body condition is critical for predicting responses to environmental change and to effective fishery management; yet low explanatory power of covariates on individual condition constitutes a major challenge.

## Introduction

Body condition is a morphometric index that describes the “plumpness” of an organism, or its weight relative to its length (Nash *et al*., 2006; Thorson, 2015). It is related to food intake rates and metabolic activity, and is often positively associated with fitness (Morgan *et al*., 2010; Thorson, 2015). In fishes, individuals with high condition have greater reproductive potential and success (Hislop *et al*., 1978; Marshall and Frank, 1999), and poor condition increases the likelihood of skipped spawning (Jørgensen *et al*., 2006; Mion *et al*., 2018) and can lower chances of survival (Dutil and Lambert, 2000; Casini *et al*., 2016b). Hence, body condition constitutes a valuable index for evaluating changes in productivity of fish stocks from ecosystem changes (Thorson, 2015; Grüss *et al*., 2020).

Because of the link to food consumption, interannual variation in condition is often associated with changes in the strength of competition for food, via changes in density of the population, competitors, or prey species (Cardinale and Arrhenius, 2000; Casini *et al*., 2006; Thorson, 2015; Grüss *et al*., 2020). Condition has also been linked to abiotic environmental variables (e.g., temperature, salinity) affecting ecosystem productivity and local habitat quality (Möllmann *et al*., 2003; Morgan *et al*., 2010; Thorson, 2015; Grüss *et al*., 2020). More recently, studies have found a link between declining body condition and deoxygenation (often resulting in the expansion of “dead zones” causing habitat degradation and compression) (Casini *et al*.,2016a, 2021), fueled by warming and nutrient enrichment (Diaz, 2001; Breitburg, 2002; Diaz and Rosenberg, 2008; Carstensen *et al*., 2014). However, reduced oxygen concentrations also cause lower food intake rates due to lower metabolic rates, which can occur even during milder hypoxia. As both environmental and biological variables can affect condition, it is important to study their relative contribution to condition in a common framework.

The Baltic Sea constitutes an interesting case study for disentangling ecosystem drivers affecting body condition (Reusch *et al*., 2018). First, in the Eastern Baltic Sea cod stock (hereafter referred to as cod), the average body growth and body condition has declined since the collapse of the stock in the early 1990s (Casini *et al*., 2016a; Mion *et al*., 2021). This has compromised the stock’s productivity to the extent that population biomass is expected to remain below safe limits despite the ban of targeted cod fisheries in 2019 (ICES, 2021a, 2021b). Second, the Baltic ecosystem has seen a major change in the abundance and distribution of both cod and its potential competitors for the benthic prey *Saduria entomon* (Haase *et al*., 2020; Neuenfeldt *et al*., 2020)—the flounder species (European flounder *Platichthys flesus* and Baltic Flounder *Platichthys solemdali*) (Orio *et al*., 2017), and its main pelagic prey species (sprat *Sprattus sprattus* and herring *Clupea harengus*) (Casini *et al*., 2011; Eero *et al*., 2012; ICES, 2021a). Also increased intraspecific competition has been linked to the low growth rates of the stock (Svedäng and Hornborg, 2014). Lastly, the irregular inflows of saline and oxygenated water from the North Sea together combined with a slow water exchange (a residence time of 25–30 years) are features that have contributed to making the Baltic Sea the largest anthropogenically induced hypoxic area in the world (Carstensen *et al*., 2014). It is also one of the fastest warming regional seas (Belkin, 2009; Reusch *et al*., 2018). Previous studies have linked changes in mean condition of cod over large spatial scales to single or some combination of ecosystem drivers (Casini *et al*., 2016a, 2021; Orio *et al*., 2020). However, in previous studies, within-population variability in condition have been neglected and the effects of environmental and biotic covariates have not been studied on local scales. Moreover, the effect of all the above-mentioned covariates on cod condition have not been analyzed in a common framework.

In this study, we apply geostatistical models to characterize the spatiotemporal variation in individual body condition and distribution of cod in the south-eastern Baltic Sea. We use data from the scientific surveys between 1993–2019, which corresponds to a period of initially high but then deteriorating cod condition (Casini *et al*., 2016a). We then seek to (1) identify which set of covariates (biomass densities of flounder and cod (representing competition), *S. entomon* (benthic prey), biomass of sprat and herring (pelagic prey), as well as depth, oxygen concentration and temperature), at different scales, can explain observed variation in condition; and (2) explore the role of changes in the spatiotemporal distribution of cod in observed trends in body condition.

## Materials and methods

### Data

To model the spatiotemporal development of cod condition and distribution, we acquired weight and length data, as well as catch per unit effort data (CPUE, numbers/hour) of cod by 10-mm length class from the Baltic International Trawl Survey (BITS) between the years 1993–2019 in the International Council for the Exploration of the Sea (ICES) subdivisions 24–28 (*SI Appendix*, Fig. S1). CPUE data were standardized based on gear dimensions and towing speed following Orio *et al*. (2017) to the unit kg/km^2^ using a TVL trawl with 75 m sweeps (note that compared to Orio *et al*. (2017), we further express density in kg/km^2^ instead of kg in 1 h trawling, sweeping an area of 0.45 kg/km^2^ by dividing by 0.45). Abundance density was converted to biomass density by fitting annual weight-length regressions. We used only data from the fourth quarter (mid-October to mid-December), which corresponds to the main growing and feeding season of cod (Aro, 1989) and also the quarter in which the Baltic International Acoustic Survey (BIAS) is conducted, meaning sprat and herring biomass can be used as covariates. The BITS data can be downloaded from https://www.ices.dk/data/data-portals/Pages/DATRAS.aspx.

### Estimating spatiotemporal development of body condition and biomass density

#### Condition model

We modelled cod condition using a spatiotemporal version of Le Cren’s relative condition factor (*K_rel_*). This factor is defined as the ratio between the observed weight for individual fish *i*, caught in time *t* at space *s*, and the predicted weight. The predicted weight was given by the relationship 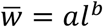, where parameters *a* and *b* were estimated in a non-spatial model with all years pooled, to represent the average weight prediction, 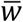 based on observed lengths *l*. An individual cod with a *K_rel_* = 1 thus has the average condition across years and space. Unlike Fulton’s K, Le Cren’s relative condition factor does not rely on the assumption that growth is isometric (*b* = 3), which, if violated, leads to bias when comparing condition of different lengths as the condition factor scales in proportion to *l*^*b*–3^ (Le Cren, 1951). Spatially correlated residual variation was accounted for with spatial random effects through Gaussian random fields. This approach to modelling spatiotemporal data is an increasingly popular method for explicitly accounting for spatial and spatiotemporal variation due to its ability to improve predictions of fish density (Thorson *et al*., 2015a) and range shifts (Thorson *et al*., 2015b), and its availability in open source software such as the R package ‘INLA’ (Rue *et al*., 2009; Lindgren *et al*., 2011).

To assess the ability of covariates (see section *Covariates* below) to explain variation in condition, we fit a geostatistical generalized linear mixed-effects model (GLMM) to the natural log of spatiotemporal Le Cren factor, assuming Student-t distributed residuals (with the degrees of freedom parameter [*ν*] set to 5) due to the presence of extreme values:

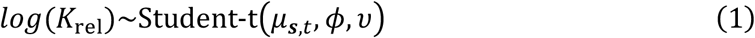

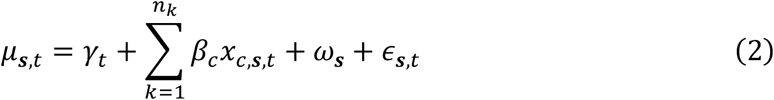

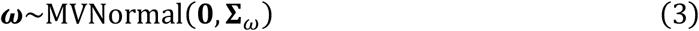

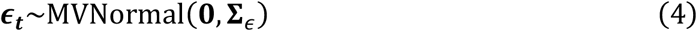

where *K_rel_* represents the Le Cren condition factor at space ***s*** (a vector of two UTM zone 33 coordinates) and time *t*, *μ* represents the mean weight, and *ϕ* represents the scale parameter. The parameters *γ_t_* represent independent means for each year. The variable *x_c,s,t_* represents the *c*-th covariate (biomass densities of flounder and cod, biomass of sprat, herring, and *S. entomon*, depth, oxygen concentration and temperature) and *ß_c_* is the covariate’s effect. The parameters *ω_s_* and *ϵ_s,t_* (Eq. 3-4) represent spatial and spatiotemporal random effects, respectively. Spatial and spatiotemporal random effects were assumed to be drawn from Gaussian random fields (Lindgren *et al*., 2011; Cressie and Wikle, 2015) with covariance matrices **∑**_*ω*_ and **∑**_*ϵ*_. The covariance (Φ(*s_j_*, *s_k_*)) between spatial points *S_j_* and *s_k_* in all random fields is given by a Matérn function.

#### Density models

We fit spatiotemporal GLMMs to biomass density data in a similar fashion as for condition to 1) evaluate how the depth distribution of cod, as well as oxygen and temperature conditions experienced by cod, have changed; and 2) use predicted local densities of cod and flounder as covariates in the condition model. For the first task, we used the predicted density at space s and time *t* as weights when calculating the annual median (and interquartile range) depth, temperature, and oxygen concentration.

We modelled densities using a Tweedie distribution, as density is both continuous and contains 0 values (Tweedie, 1984; Shono, 2008; Anderson *et al*., 2019):

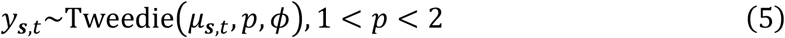

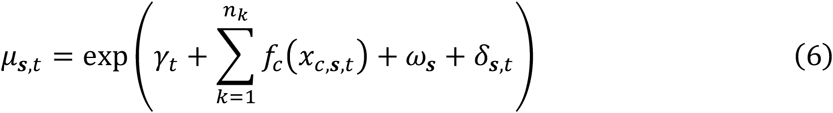

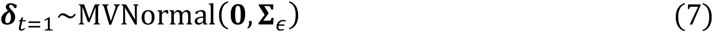

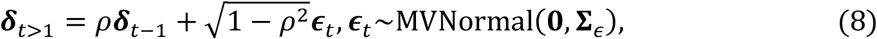

where *y_s,t_* represents density (kg/km^2^) at space ***s*** and time *t*, *μ* is the mean density, and *p* and *ϕ* represent power and dispersion parameters, respectively. The parameters *γ_t_* represent independent means for each year, *f_c_* is a penalized smooth function for covariate *x_c_*, and *ϵ_s,t_* represent spatial and spatiotemporal random effects. The parameters *ω_s_* and have the same definition as in the condition model (Eq. 3), but the spatiotemporal random effects are here assumed to follow a stationary AR1-process where *ρ* represents the correlation between subsequent spatiotemporal random fields.

#### Covariates

For both models (condition and density), covariates were chosen to reflect hypothesized drivers based on published literature. For the condition model, we included covariates at spatial scales that roughly reflect the habitats cod would have been exposed to during the seasonal build-up of energy reserves. Recent tagging studies suggest cod are either stationary or mobile over the course of a year moving between feeding and spawning habitats (Mion *et al*., 2022). However, within the feeding season, cod move roughly over an area corresponding to an ICES rectangle (1° by 30’, *SI Appendix* Fig. S1) (Hüssy *et al*., 2020). Therefore, we included environmental and biological demersal covariates (sea bottom temperature [°C], sea bottom oxygen [ml/L], depth [m], and biomass density of cod and flounder [kg/km^2^] and *S. entomon* [mg/m^2^]) at the haul level and the median over the ICES rectangle-level. The pelagic covariates were included at the ICES rectangle- and subdivision-level (as pelagic species are highly mobile) (see *SI Appendix*, Fig. S1 for the spatial units ICES rectangle and subdivision).

Biomass of sprat and herring (tones) were extracted from the ICES WGBIFS database for the BIAS survey data (https://www.ices.dk/community/groups/pages/WGBIFS.aspx). Monthly predictions for sea bottom temperature and sea bottom concentration of dissolved oxygen were extracted at the haul locations from the ocean model NEMO-Nordic-SCOBI (Eilola *et al*., 2009; Almroth-Rosell *et al*., 2011; Hordoir *et al*., 2019) and averaged for October–December (approximately 14%, 76% and 10% of the BITS hauls were conducted in October, November and December, respectively). We also conducted preliminary analysis to determine if oxygen should be modelled with a linear (as depicted in Eq. 6), or a linear threshold effect, as suggested in experimental studies (Chabot and Dutil, 1999; Hrycik *et al*., 2017). This showed that the model with a linear effect was favored in terms of Akaike Information Criterion (AIC) (*SI Appendix*, Table S1). Depth raster files were made available by the EMODnet Bathymetry project, https://www.emodnet.eu/en/bathymetry, funded by the European Commission Directorate General for Maritime Affairs and Fisheries. Biomass density of *S. entomon* was extracted from a habitat distribution model coupled with modelled hydrographical data from the regional coupled ocean biogeochemical model ERGOM (Gogina *et al*., 2020; Neumann *et al*., 2021). We used predicted densities of cod and flounder (kg/km^2^) from GLMMs (described above) as covariates, since not all hauls in the CPUE (density) data could be standardized and joined with the condition data. For the cod and flounder models that were used to provide covariates for the condition model, the only covariate used was depth. For the cod density models used to evaluate effects of changes in the average depth, oxygen concentration and temperature, we used only these three variables and a fixed year effect as covariates.

Following Thorson (2015) and Grüss *et al*. (2020), we rescaled all covariates to have a mean of 0 and a standard deviation of 1. This facilitates comparison between covariates of different units and allows for comparison between the estimated coefficients and the marginal standard deviation of spatial (*σ_0_*) and spatiotemporal (*σ_E_*) variation. We did not conduct any model selection after our *a priori* selection of covariates to avoid statistical issues with inference from stepwise selection (e.g., Whittingham *et al*., 2006), and because initial analyses suggested the model was not overfit (see *SI Appendix*, Fig. S2 for Pearson correlation coefficients across variables).

#### Model fitting

For computational efficiency, we fit all models in a “predictive process” modelling framework (Latimer *et al*., 2009; Anderson and Ward, 2019), where spatial and spatiotemporal random fields are approximated using a triangulated mesh and the SPDE approximation (Lindgren *et al*., 2011) (*SI Appendix*, Fig. S3, S12), created using the R-package ‘R-INLA’ (Rue *et al*.,2009). The random effects were estimated at the vertices (“knots”) of this mesh and bilinearly interpolated to the data locations. The locations of the knots were chosen using a *k*-means clustering algorithm, which minimizes the total distance between data points and knots. As the knot random effects are projected to the locations of the observations, more knots generally increase accuracy at the cost of computational time. After initial exploration, we chose 100 knots for the condition model and 200 knots for the density models. We fit the models using ‘TMB’ (Kristensen *et al*., 2016) via the R-package ‘sdmTMB’ (version 0.1.0) (Anderson *et al*.,2022) with maximum marginal likelihood and the Laplace approximation to integrate over random effects. We assessed convergence by confirming that the maximum absolute gradient with respect to all fixed effects was < 0.001 and that the Hessian matrix was positive-definite. Model residuals are shown the *SI Appendix* (Figs. S4-S6 and S13-S14). We used packages in the ‘tidyverse’ (Wickham *et al*., 2019) for data processing and plotting.

## Results

The condition model revealed a mean decline in the Le Cren condition factor of 17% [11%, 20%] (values in brackets are the 2.5% and 97.5% quantiles from 500 draws from the joint precision matrix). The condition factor declined from approximately 1.15 to 0.95 between 1993 and 2019 (the decline leveled off around 2008) and the whole distribution of condition values in the population exhibited a shift to lower values (Fig. 1A). The condition factor declined the most in the northern subdivisions (i.e., 27 and 28, where also the biomass of cod is lowest, Fig. 4A) and the least in the south-western subdivision 24 (Fig. 1B). The spatial predictions from the condition model illustrate the presence of consistent “low spots” of body condition in deep and low-oxygen areas, and that the condition factor declined in the whole area over time (Fig. 2, *SI Appendix*, Fig. S9).

**Fig. 1.**
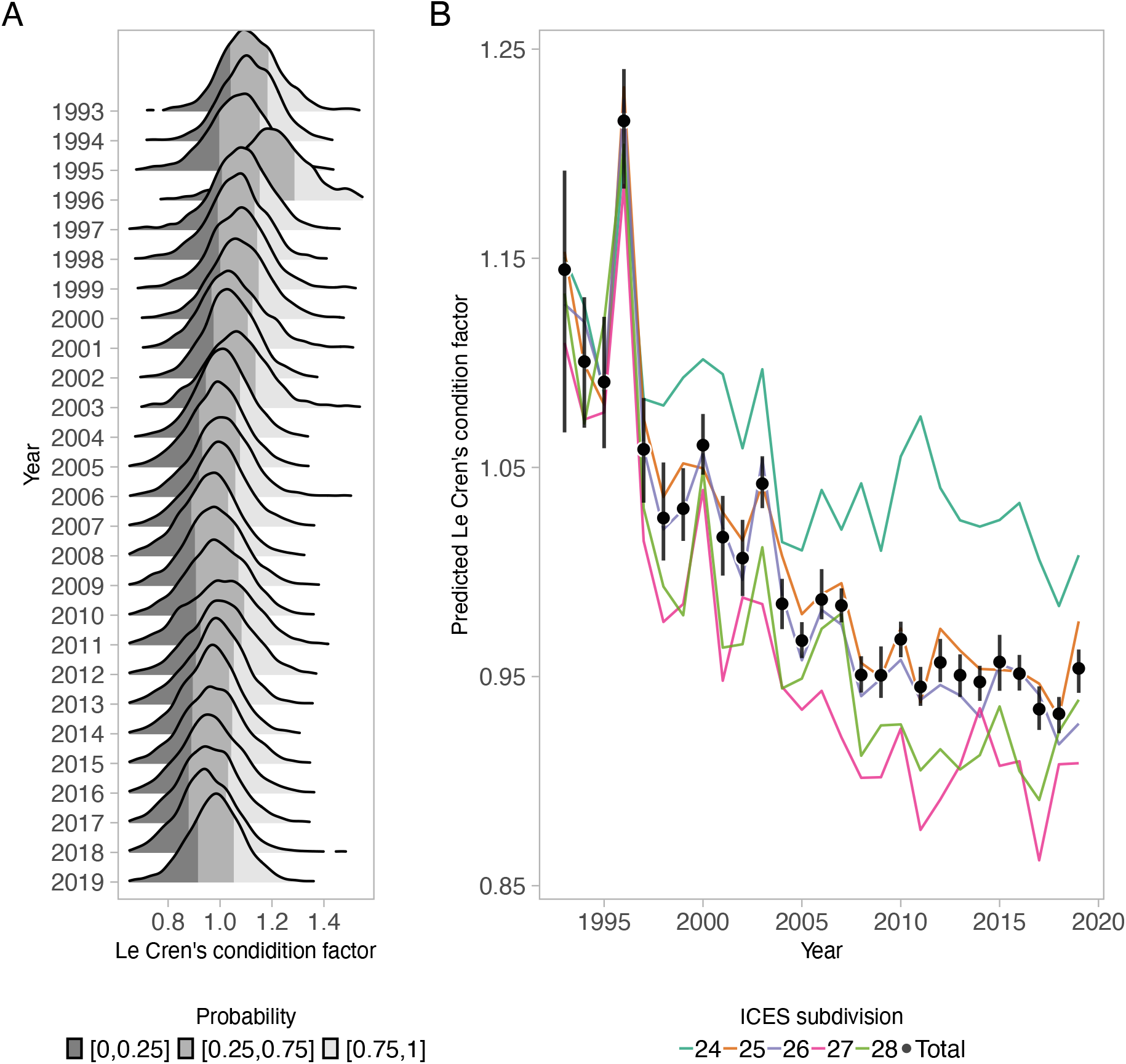
A) Density plots of condition data by year. B) Predicted Le Cren condition factor of cod over the period 1993-2019 in the Baltic Sea (total as well as by each ICES subdivisions), acquired by predicting from the spatiotemporal condition model over a grid with spatially varying covariates set to their true values (ICES rectangles with missing pelagic data were given the subdivision median when predicting but not fitting, see *SI Appendix*, Fig. S23). Vertical line segments depict 95% confidence intervals.

**Fig. 2.**
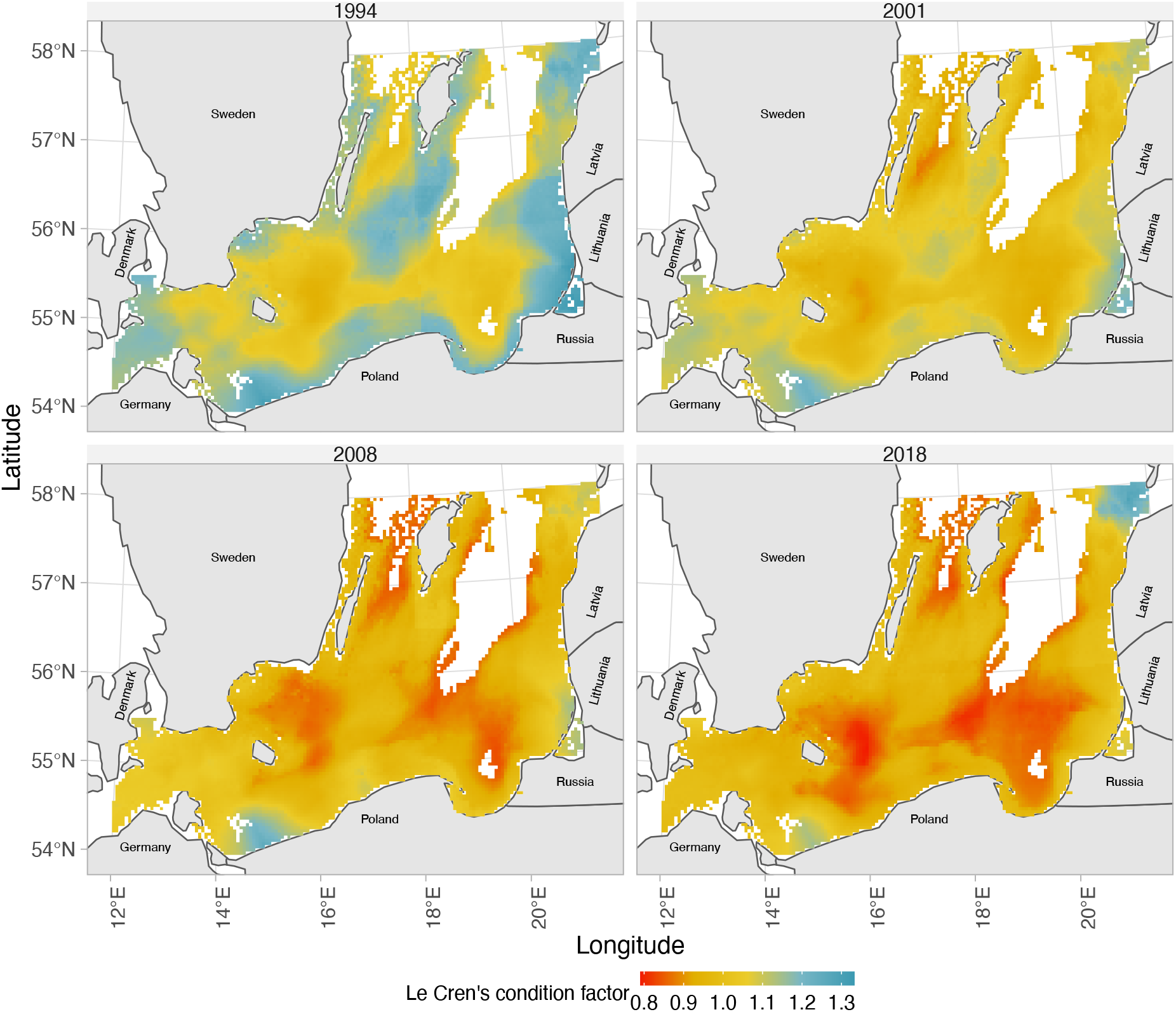
Predicted Le Cren’s condition factor with spatially varying covariates set to their true values (ICES rectangles with missing pelagic data were given the subdivision mean, see *SI Appendix*, Fig. S23). Included in the plot are years 1994, 2001, 2008, 2018. Only grid cells with depths between 10m and 120m are included in the plot. For all years in the series, see *SI Appendix*, Fig. S9.

The covariates with the largest positive standardized effect sizes on the condition factor were median depth of the ICES rectangle (0.006 [0.001–0.012]), temperature at the haul level (0.008 [0.004–0.011]) (values in brackets indicate 95% confidence interval), biomass of sprat at the ICES subdivision level (0.012 [0.006, 0.018]) and oxygen concentration at the ICES rectangle level (0.007 [0.0014, 0.0124]) (Fig. 3). Temperature and biomass density of *S. entomon* at the rectangle level and oxygen at the haul level had smaller positive effects. Depth at the haul level was negatively associated with condition (−0.025 [−0.028, −0.022]) (see *SI Appendix*, Fig. S10, for conditional effects plots), and so was cod density at haul and rectangle level (−0.0014 [−0.0038, 0.001] and −0.0009 [−0.0046, 0.0028], respectively). The biomass density of *S. entomon* at the haul level and the biomass of sprat at the rectangle level, as well as the density of flounder and the biomass of herring at any scale, had smaller and more uncertain effects on condition (Fig. 3). The effect sizes of fixed effects were several times smaller than the magnitude of latent spatiotemporal and spatial variation (Fig. 3). Using the approach proposed in Nakagawa and Schielzeth (2013), we calculated the marginal *R*^2^ for fixed and random effects, and found that fixed effects had a marginal *R*^2^ of 0.153 (0.1 for fixed year effects and 0.06 for the remaining covariates), while random effects had a marginal *R*^2^ of 0.218 (0.08 for spatial random effects and 0.13 for spatiotemporal random effects).

**Fig. 3.**
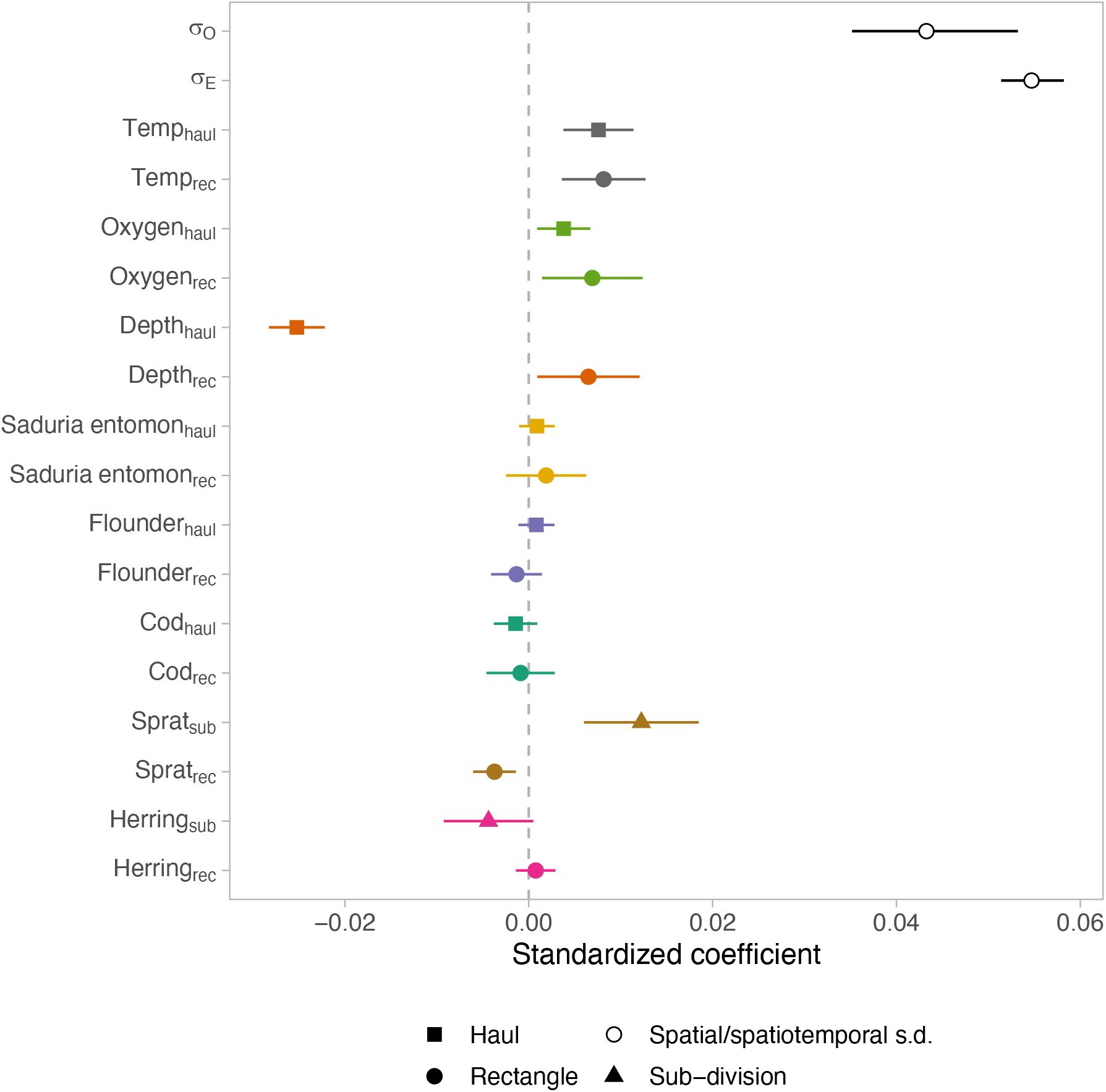
Mean and 95% confidence interval of standardized coefficients (effect sizes) for covariates and spatial and spatiotemporal standard deviation (*σ_E_* and *σ*_0_, respectively) in the condition model. The subscript haul refers to covariates estimated at the location of the haul, rec refers to covariates at the ICES statistical rectangle and sub refers to covariates over ICES subdivisions (*SI Appendix*, Fig. S1). Colors indicate covariate-groups and shapes indicate scale.

**Fig. 4.**
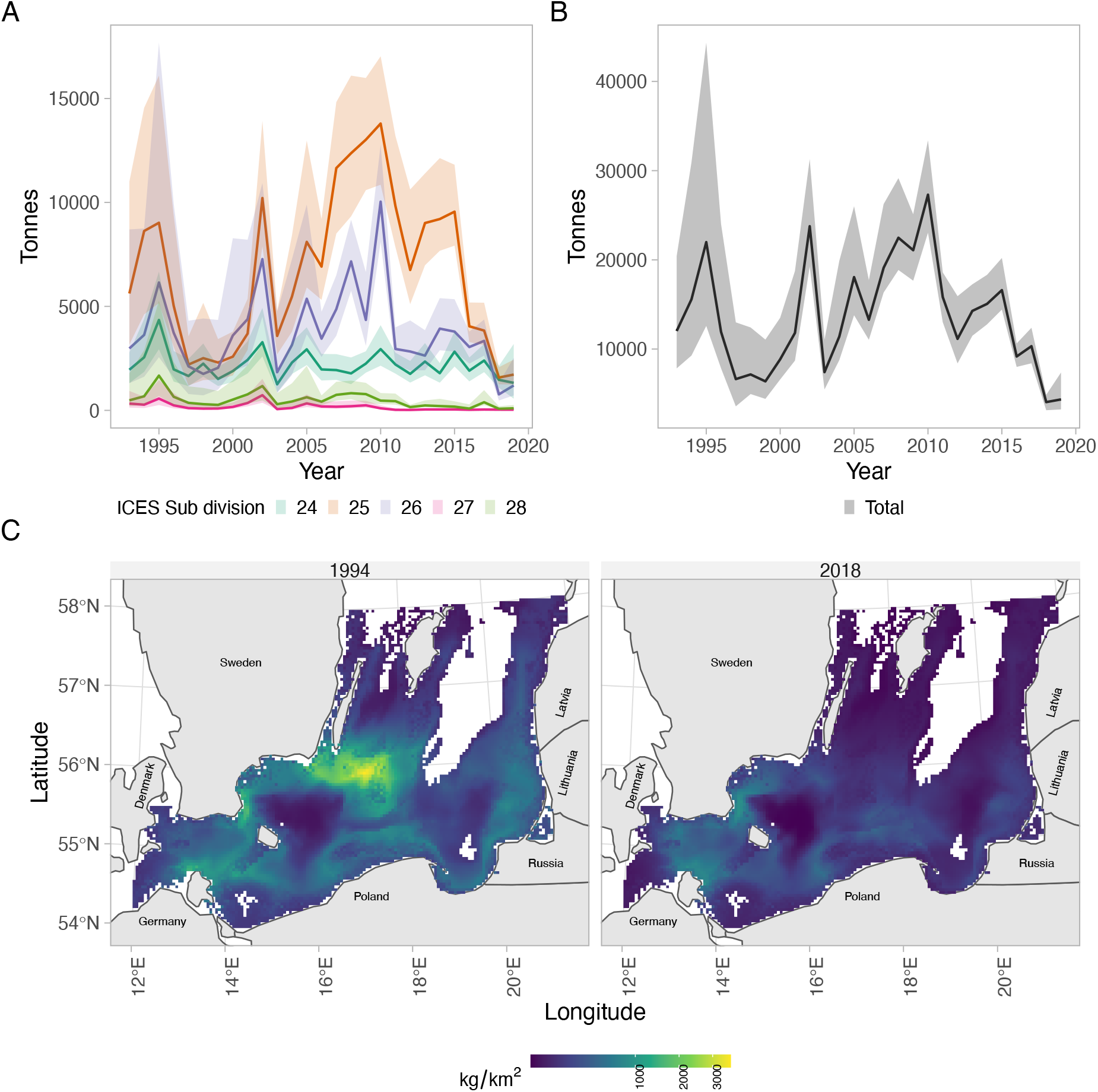
A) Predicted biomass from the spatiotemporal density model by ICES subdivision (B) and total across all subdivisions. C) Predicted density (kg/km^2^) in select years (1994 and 2018). For all years in the series, see *SI Appendix*, Fig. S17. Only grid cells with depths between 10 and 120 are included in the plot.

We conducted a sensitivity analysis by fitting the condition model to different parts of the data. The different models were only cod above 30 cm, only cod below 30 cm, omitting subdivision 24 (the mixing zone with western Baltic cod (Mion *et al*., 2022)), and including only grid-points with cod above a certain threshold when calculating median variables across the ICES rectangle. However, the model coefficients were similar across all models (*SI Appendix*, Fig. S11).

The median depth and oxygen experienced by cod (depth and oxygen weighted by the predicted biomass density of cod at location, respectively, Fig. 4C) got deeper and declined, respectively, throughout the time period (Fig. 5). However, the population again occupied slightly shallower waters in the last 3 years of the time series (Fig. 5C; see *SI Appendix*, Fig. S21 for results split by subdivision, Fig. S24 for the corresponding analysis on temperature). The trends in experienced oxygen were steeper than the average oxygen in the environment at depths corresponding to the interquartile range of cod (Fig. 5C-D). In fact, the average oxygen concentration in the environment declined by approximately 0.65 ml/L between 1993 and the lowest in 2006, while the biomass-weighted oxygen concentration declined more steadily (approximately 1 ml/L between 1993 and 2019) (*SI Appendix*, Fig. S19-S20 for estimates split by subdivision). However, while the biomass-weighted oxygen concentration declined between 1993 and 2019) (Fig. 5D), the corresponding effect on condition given the effect size of oxygen at the haul was small (*SI Appendix*, Fig. S10). The standardized effect size for oxygen of 0.004 [0.0009, 0.0067] means that for each unit increase in the variable (i.e., 1 standard deviation or 1.85 ml/L), the Le Cren condition factor increased by 0.4%. This can be compared to the 1 ml/L decline in the oxygen concentration and 17% decline in the condition factor between 1993–2019.

**Fig. 5.**
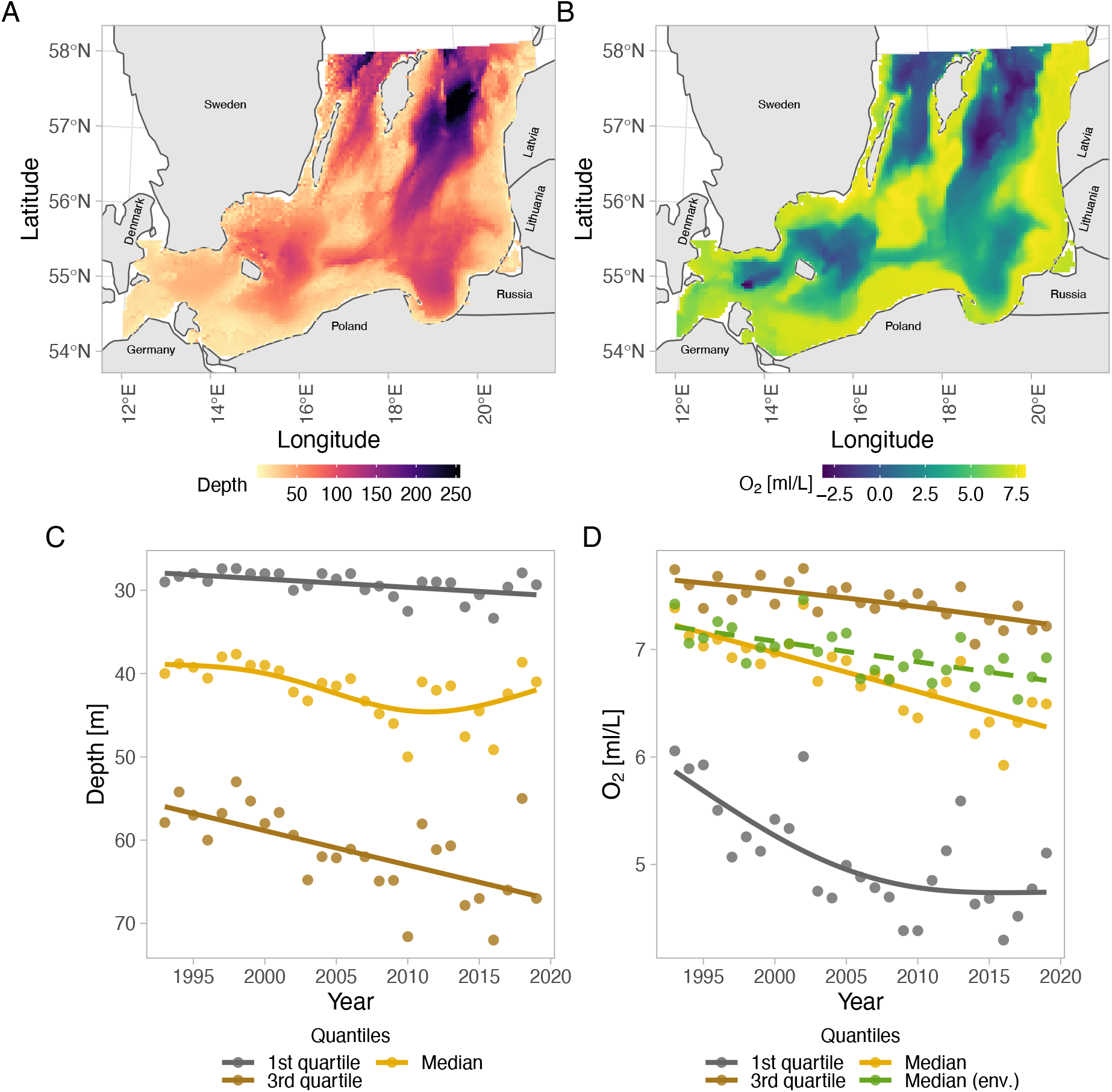
(A) Bathymetry and (B) oxygen concentration (exemplified using year 2006) in the study area. Panels (C) and (D) illustrate depth and oxygen weighted by predicted cod density, respectively. Colors indicate biomass-weighted quantiles (1^st^ quartile, median and 3^rd^ quartile), as well as the un-weighted average (green dashed line) at depths corresponding to the average interquartile range (29–61 m). Lines depict generalized additive model (GAM) fits (*k*=4).

## Discussion

The body condition of fish depends on previous energy accumulation and is therefore largely shaped by the quality of the habitat the fish has occupied. By using a spatially explicit condition model, we can link the condition of Eastern Baltic cod to covariates at different ecologically relevant spatial scales. Our model reveals that the Le Cren condition factor declined on average by 17%, in 1993–2019, with most of this decline occurring 1993–2008. Moreover, while there are persistent low-spots of body condition (in the deep and low-oxygen areas), the condition declined in the whole area, which suggests that there are drivers acting on large spatial scales. While we identify changes in the spatiotemporal distribution of cod that could have led to poorer environments experienced by cod (deeper areas with less oxygen), effect sizes of single covariates were overall small, while latent spatial and spatiotemporal variation was several times larger in magnitude and explained more variation in condition. One likely explanation for this is that we use individual-level body condition, while previous studies using average condition as a response variable (but the same large-scale covariates) generally find stronger relationships (Casini *et al*., 2016a).

Previous studies have suggested both direct (Limburg and Casini, 2019; Brander, 2020) and indirect (Brander, 2020, 2022; Neuenfeldt *et al*., 2020; Orio *et al*., 2020) effects of oxygen as a cause for the declining body condition of cod in the past three decades. Direct effects here refer to mild hypoxia reducing the appetite and food consumption (Chabot and Dutil, 1999) and, by extension, also their condition, as their ability to accumulate energy reserves reduces. We found that Baltic cod experienced oxygen concentrations at around 7.2 [5.8–7.7] (interquartile range in brackets) ml/L on average (biomass-weighted median) in 1993 and are currently experiencing oxygen concentrations at around 6.3 [4.5–7.2] ml/L. In subdivision 25 (the core area of cod, currently) we estimate it to be around 6.5 [4.9–7.3] ml/L in 2019 (*SI Appendix*, Fig. S19). This is higher than recent estimates of an average oxygen concentration of 4–4.5 ml/L, based on oxygen levels at the mean depth of the cod population in the recent years (Brander, 2020; Casini *et al*., 2021).

One reason for the difference in our estimate compared to previous studies is because instead of calculating average oxygen at the mean depth of cod, we weighted the sea bottom oxygen in the environment (from the ocean model NEMO-Nordic-SCOBI) by the predicted densities from the cod density model. This approach overcomes the issue that oxygen concentrations span a large range for any given depth and avoids the assumption that cod depth occupancy is independent of oxygen concentration. Our finding that trends in weighted and unweighted oxygen differ suggests that it is important to account for species’ heterogenous distribution. This is particularly evident in subdivision 25, where the oxygen trends in the environment are more stable, as in (63), but the experienced oxygen by cod declined (*SI Appendix*, Fig. S19). Another reason for differences between previous estimates of experienced oxygen could be due to differences among oxygen models. For example, the model developed by Lehmann et al. (2002, 2014) (the “GEOMAR” model) and used in Casini et al. 2021 (Casini *et al*., 2021) and Orio et al. (Orio *et al*., 2019), results in weighted oxygen concentrations 0.5–1 ml/L lower on average, but a less steep decline than the NEMO-Nordic-SCOBI model between 1993-2016 in subdivision 25 (*SI Appendix*, Fig. S25). Also the unweighted estimates differ approximately 0.5-1 ml/L between the models at depths between 29-61 m. Hence, although explaining the differences between the models is outside the scope of this paper, care should be taken when interpreting absolute values of oxygen concentrations.

In an experiment by Chabot & Dutil (1999), 5 ml/L (converted from 73% O_2_ saturation at 10°C, 28‰, and 1013.25 hPa) was estimated to be a critical value concentration beyond which negative effects on growth and condition were observed on cod. This value is higher than a meta-analytic estimate across fishes of a 3.15 ml/L threshold, below which negative effects on fish growth occur (Hrycik *et al*., 2017). However, despite our data spanning oxygen levels above and below these values, we do not find support for a threshold in the relationship between condition and oxygen [see also ref. 15]. Instead we found a linear positive effect of oxygen, which is in agreement with previous studies showing that exposure to low-oxygen areas is associated with low condition in Baltic cod (Limburg and Casini, 2019; Casini *et al*.,2021). However, we can only speculate if the positive association is due to higher oxygen being correlated with richer habitats that feature higher food availability, if there are direct physiological impacts at a higher threshold in the wild, or if behavioral responses (e.g., movement between high and low oxygen area) essentially remove any measurable thresholds in natural systems.

An indirect effect of declining oxygen on condition is the potential amplification of intra- and interspecific competition with flounder for shared benthic prey species, such as the isopod *S. entomon*, due to habitat contraction of cod caused by the expansion of “dead zones” (Casini *et al*., 2016a, 2021; Orio *et al*., 2019; Haase *et al*., 2020). To address the potential effects of changes in intra- and inter-specific competition, we used predicted density of flounder and cod at the haul- and at the ICES rectangle-level, as well as *S. entomon* densities as covariates. We detected a negative effect of cod haul-level density on cod condition, in line with previous studies finding density-dependent effects on growth (Svedäng and Hornborg, 2014), though the effect is uncertain and minor compared to the other predictors. We did not detect an effect of flounder density at any scale. However, biomass density is not a direct measure of competition—areas with higher densities of cod and flounder could simply also have more food. It could also be because the biomass of both cod and flounder have been at relatively low levels during the past three decades from a historical perspective (Tomczak *et al*., 2022). The effect of *S. entomon* at the rectangle level was positive, but uncertain and small in magnitude. This lack of a clear effect could be due to benthic food availability not changing dramatically over the time period in the southern Baltic Sea, as shown in Svedäng et al. (Svedäng *et al*.,2022).

A reduced availability of sprat (either changes in their size-distribution or shifting distributions and thus reduced spatial overlap) has also been linked to poor growth and condition at the population level (Gårdmark *et al*., 2015; Casini *et al*., 2016a; Neuenfeldt *et al*., 2020). In our study, using spatially resolved data, we also found positive effects of sprat biomass on cod condition at the ICES subdivision level. The biomass of sprat generally declined from the levels in the early 90’s, and this decline is more accentuated in the northern subdivisions analyzed, where cod condition also declined the most. However, despite the decline, sprat has been most abundant in subdivision 28 (*SI Appendix*, Fig. S22), while cod biomass in subdivisions 27 and 28 has been low during the study period (Fig. 4), suggesting that further analyses should be made to infer whether the decline in sprat drove the decline in condition. Overall, the fact that cod condition has declined in all areas—also in areas where high abundance of prey remains—indicates that several variables and driving processes have been involved, including variables operating on a large spatial scale

In conclusion, our study illustrates fine-scale spatiotemporal development of body condition in the eastern Baltic cod, and population-level changes in depth distribution and oxygen concentrations. The small effect sizes of the single covariates, analyzed for the first time in a common framework using individual-level data, suggest that multiple factors are responsible for the observed spatiotemporal changes in cod condition during the past 25 years. However, the small effect size of the covariates found in our models might also be explained by the fact that condition is shaped over a long time period, while trawl data and correspondent environmental predictors reveal snapshots in time. We therefore argue that analysis of condition data from surveys conducted with low temporal resolution should be complemented with e.g., tagging studies (as suggested also by Thorson (2015)), or using “life-time recorders” such as otoliths as done in Limburg and Casini (Limburg and Casini, 2019), to determine mechanistic links between condition and covariates. However, the explanatory power of latent variables (spatial and spatiotemporal terms) in the models suggests that other factors, not explicitly included in our analyses, may have also played an important role in the decline of cod condition during the past three decades. It is also possible that the mechanisms that initiated the body condition decline are not the same ones that have kept cod in a poor physiological state in the last 10 years (Tomczak *et al*., 2022). Liver parasites, for instance, are numerous now that cod are in poor condition, but likely did not cause the decline as cod in good condition are not as susceptible to parasite infection (Ryberg *et al*., 2020). Evaluating factors associated with condition hotspots would help understand the role of food availability for condition. The Eastern Baltic cod stock is not predicted to grow even in the absence of fishing mortality (ICES, 2021a). This makes it crucial to understand how environment and species interactions affect the body condition of cod (Eero *et al*., 2020) since body condition is a key biological trait affecting mortality and reproductive output.

## Acknowledgements

We are very grateful for help from Alessandro Orio for standardization of survey data used in the density models, Federico Maioli for helpful modelling discussion, Hagen Radtke and Ivan Kuznetsov for assistance in acquiring predictions of *S. entomon* densities, Martin Hansson and Elin Almroth Rosell at SMHI for assistance with environmental data, and Olavi Kaljuste for providing pelagic data. We thank the staff involved in the scientific sampling and analysis of biological data. The study was financed by the Swedish Research Council Formas (grant no. 2018-00775 to M.C.).

## Author Contributions

All authors contributed to the manuscript. Specifically, M.C. coordinated the study, M.L. prepared the raw data, M.G. provided *S. entomon* data, M.L. led the design and conducted the statistical analyses with critical contribution from S.C.A and input from M.C. M.L. wrote the first draft. All authors contributed to revisions and gave final approval for publication.

## Data and code availability

All code and data are publicly available at https://github.com/maxlindmark/cod_condition and will be deposited on Zenodo upon publication.

