## Supporting information for "Evaluating drivers of spatiotemporal individual condition of a bottom-associated marine fish"

[1] Author to whom correspondence should be addressed. Current address:

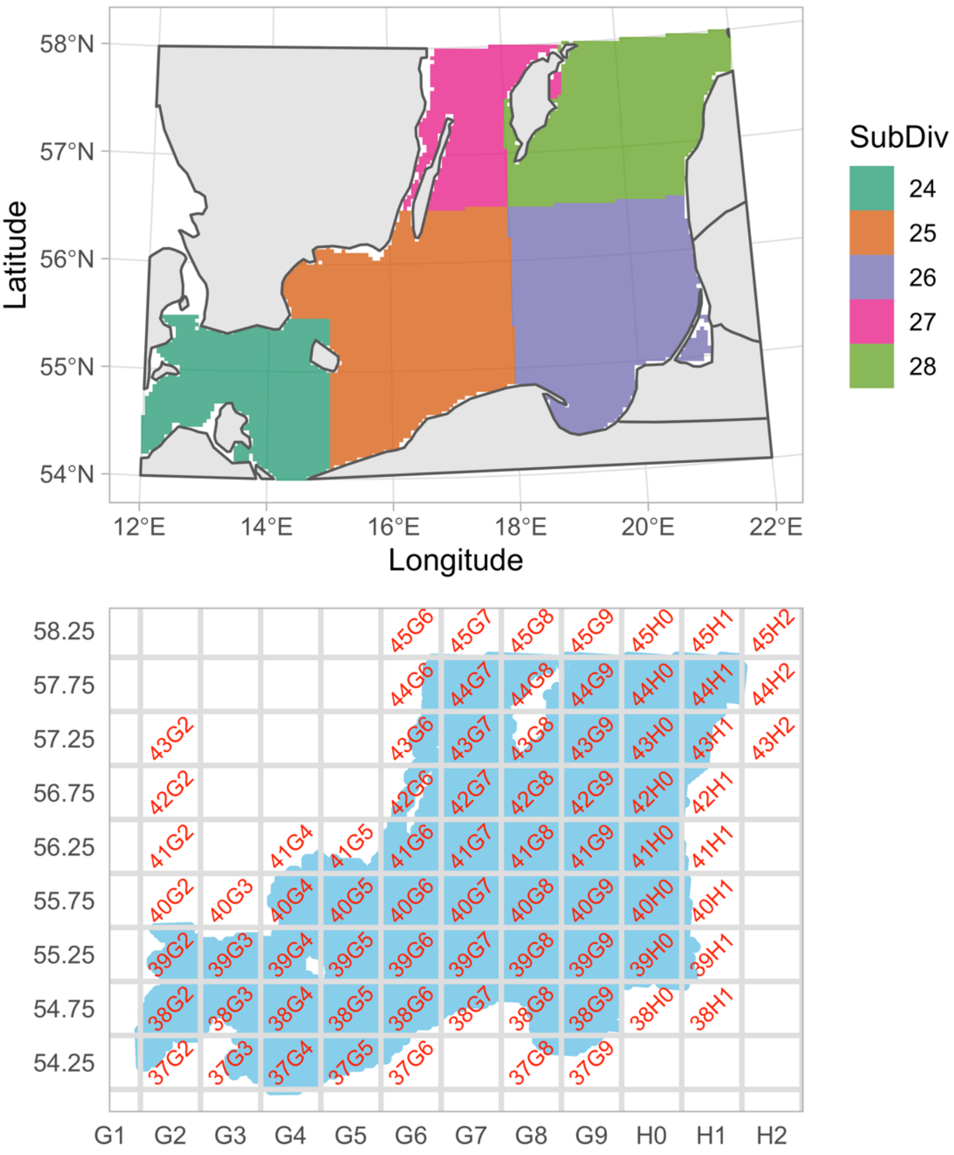


Fig. S1. Map of ICES subdivisions (top) and ICES rectangles (bottom).

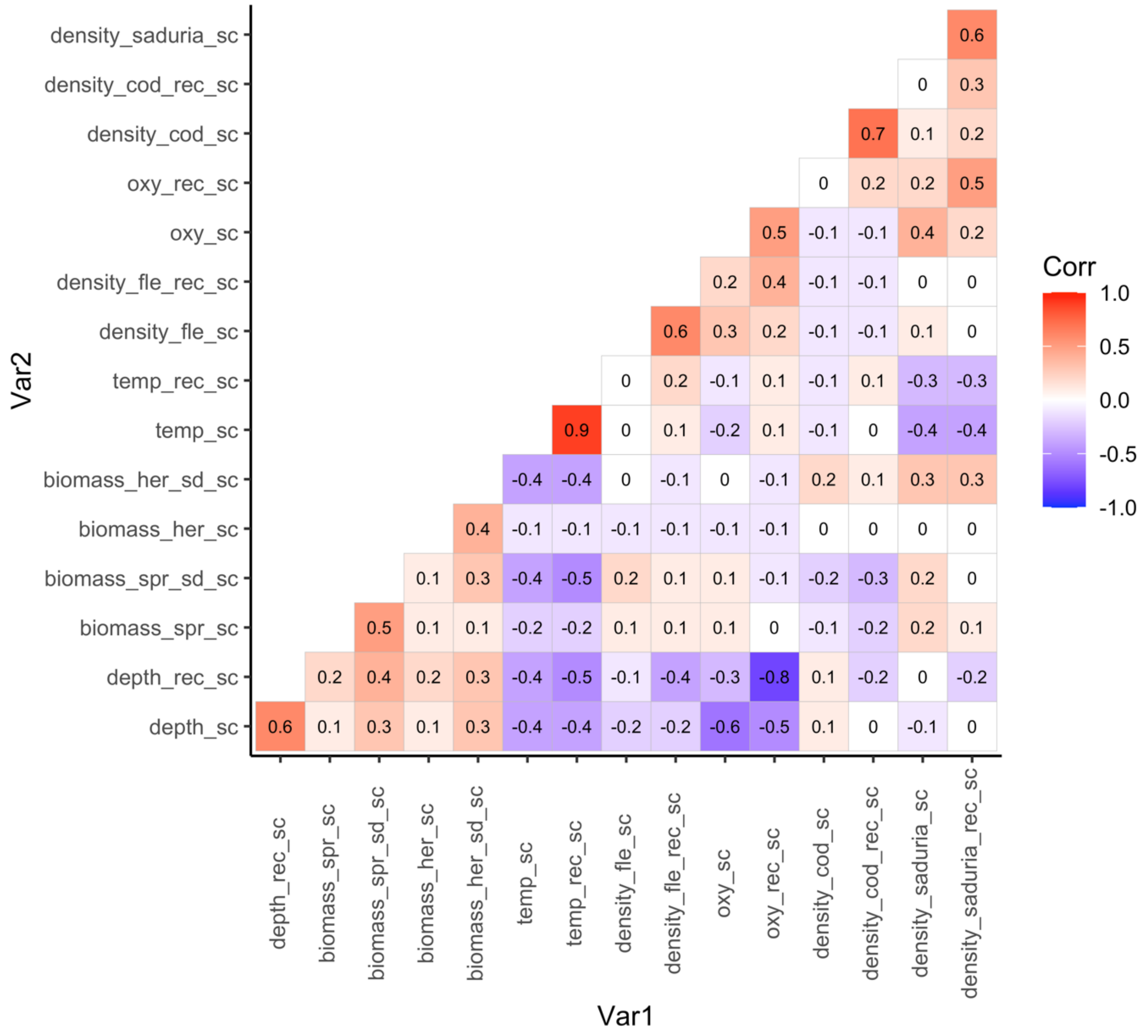


Fig. S2. Pearson correlations coefficients between all variables included in the condition model.

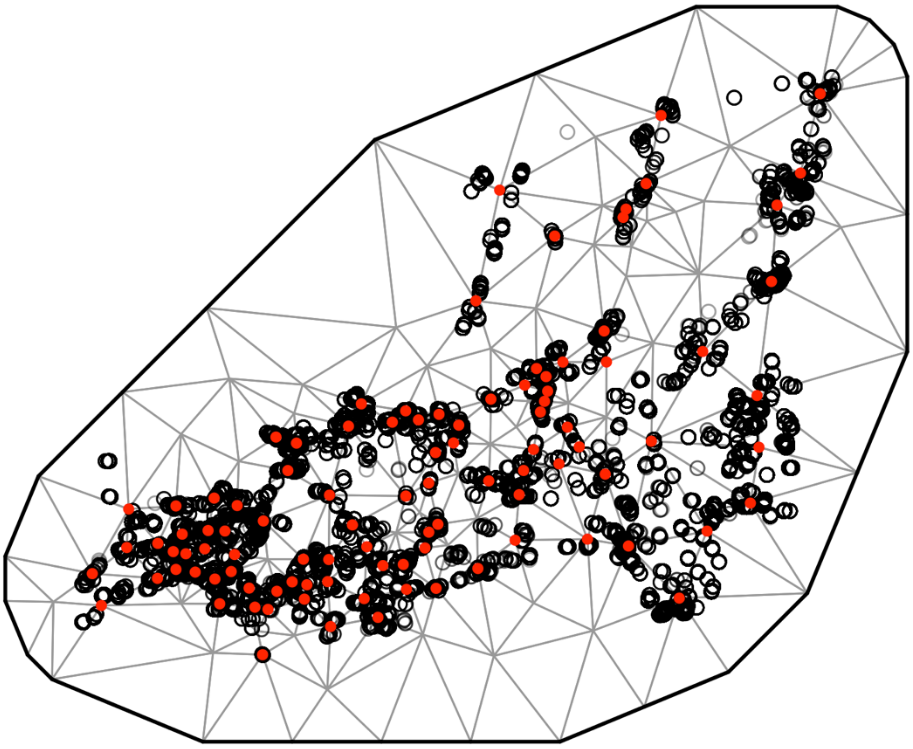

Fig. S3. SPDE mesh for condition model (100 knots).

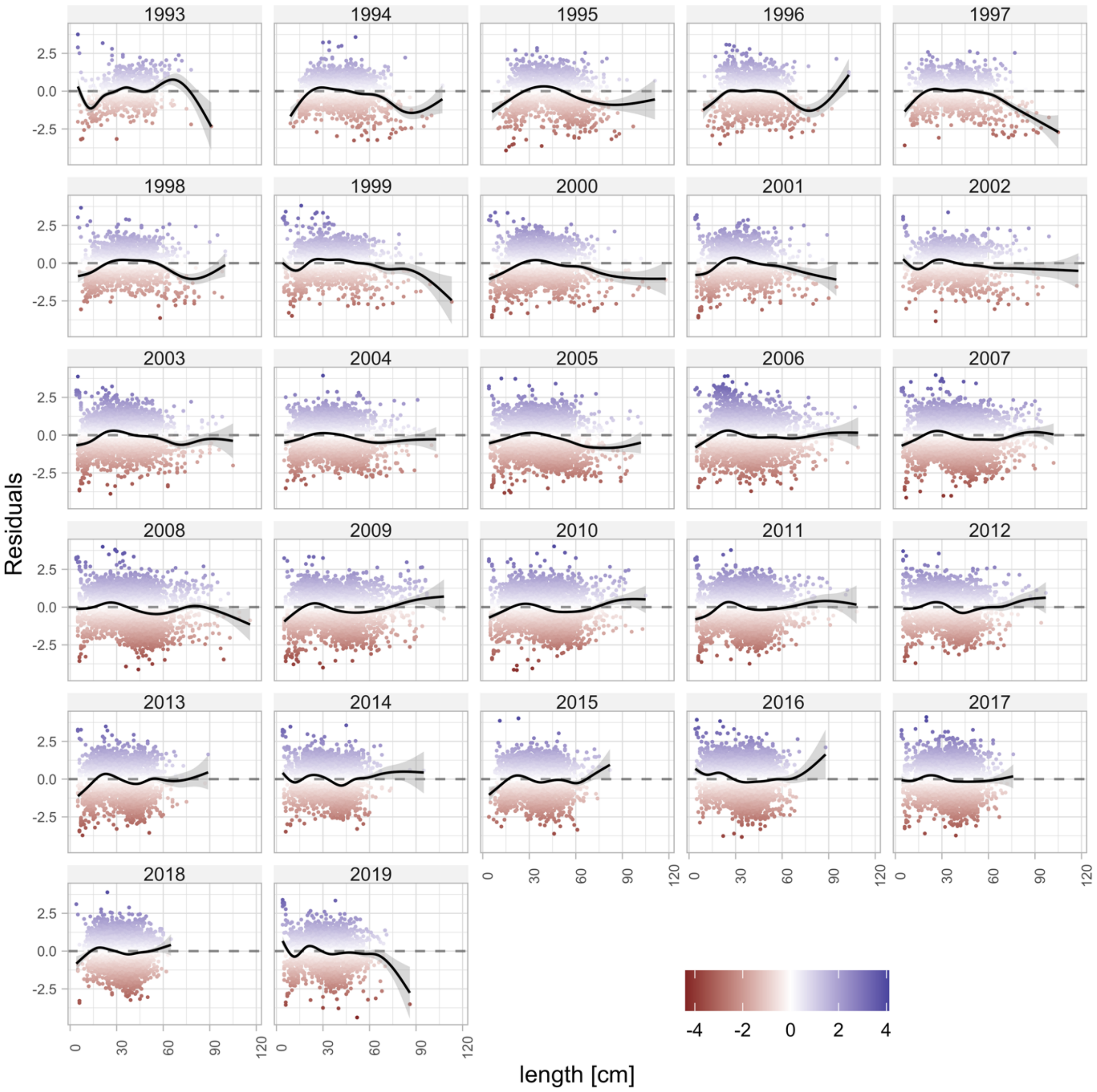


Fig. S4. Residuals from the condition model plotted against length for each year.

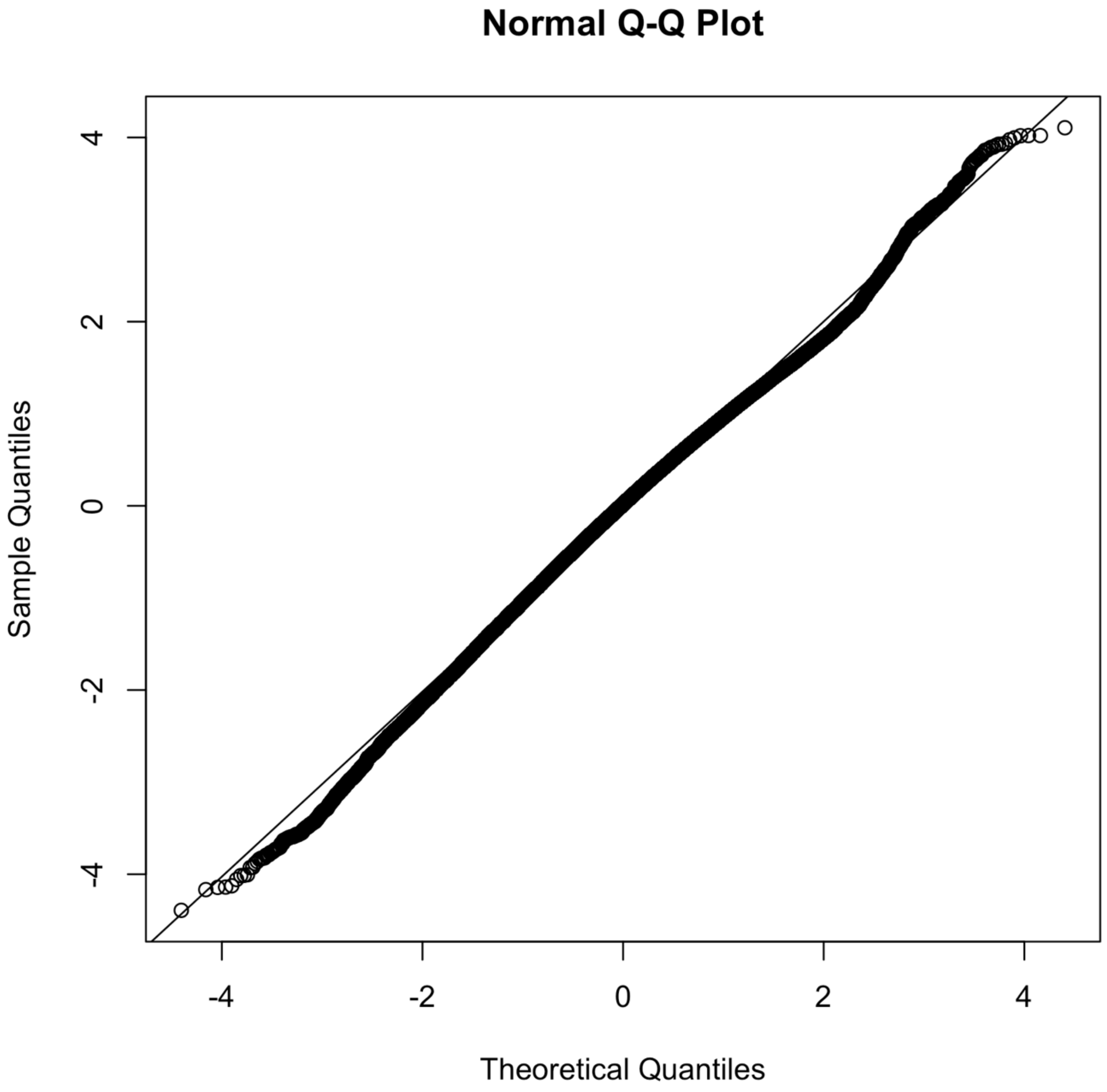


Fig. S5. QQ plot of condition model based on randomized quantile residuals (fixed effects held at their maximum likelihood estimate and the random effects are sampled with MCMC via tmbstan [1] and Stan [2], in line with [3,4].

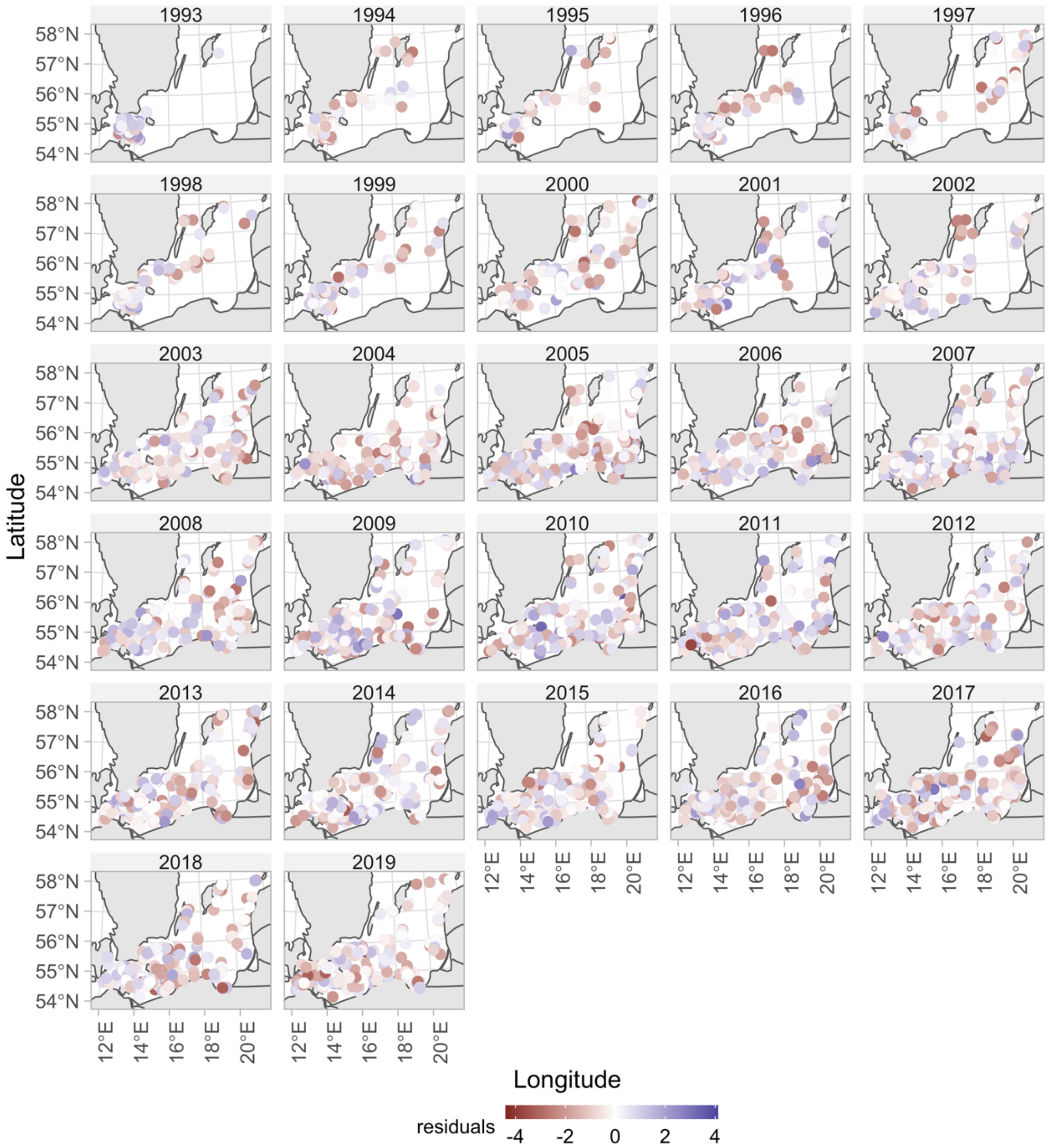


Fig. S6. Condition model residuals plotted in space.

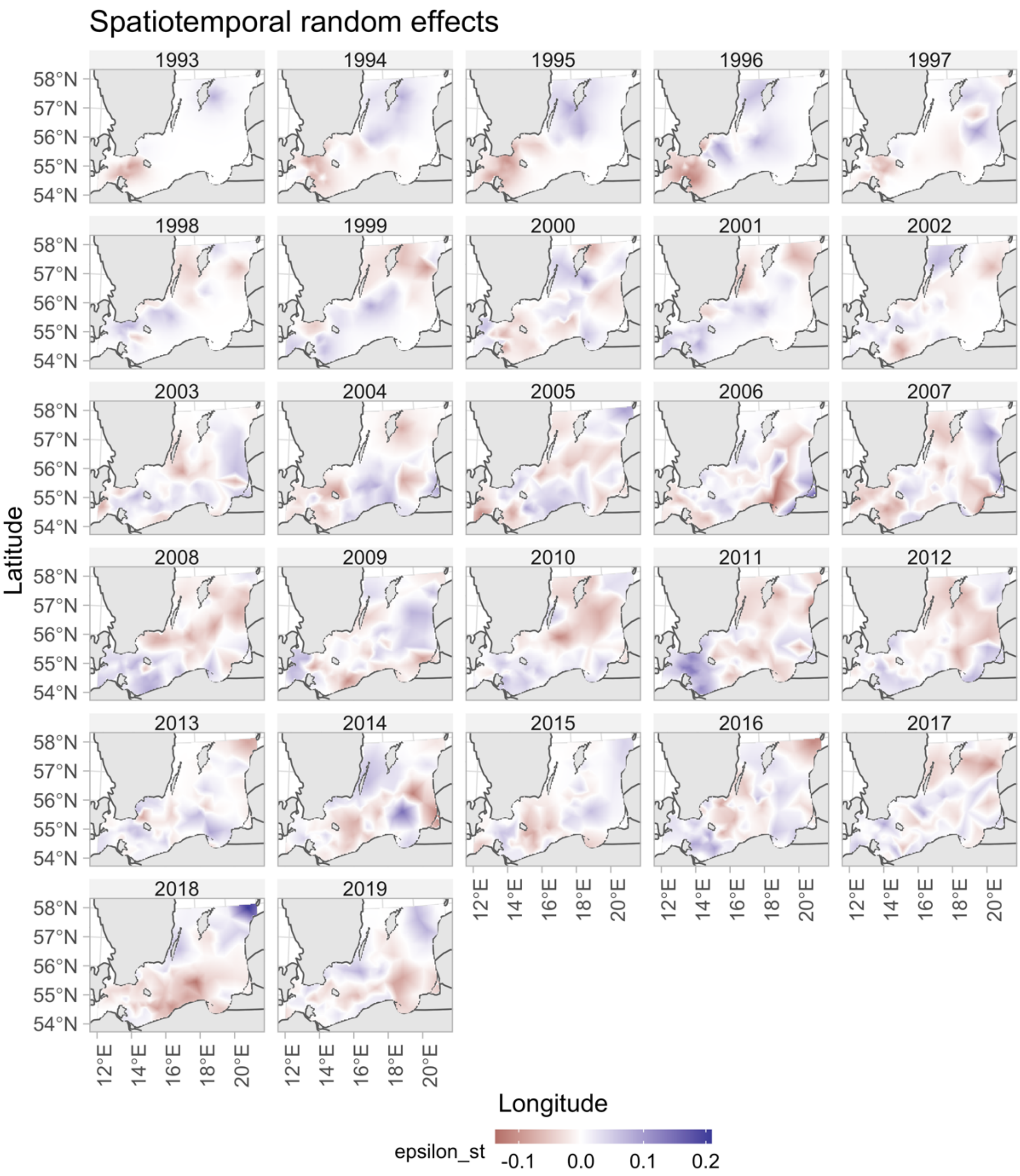


Fig. S7. Spatiotemporal random effects for the condition model.

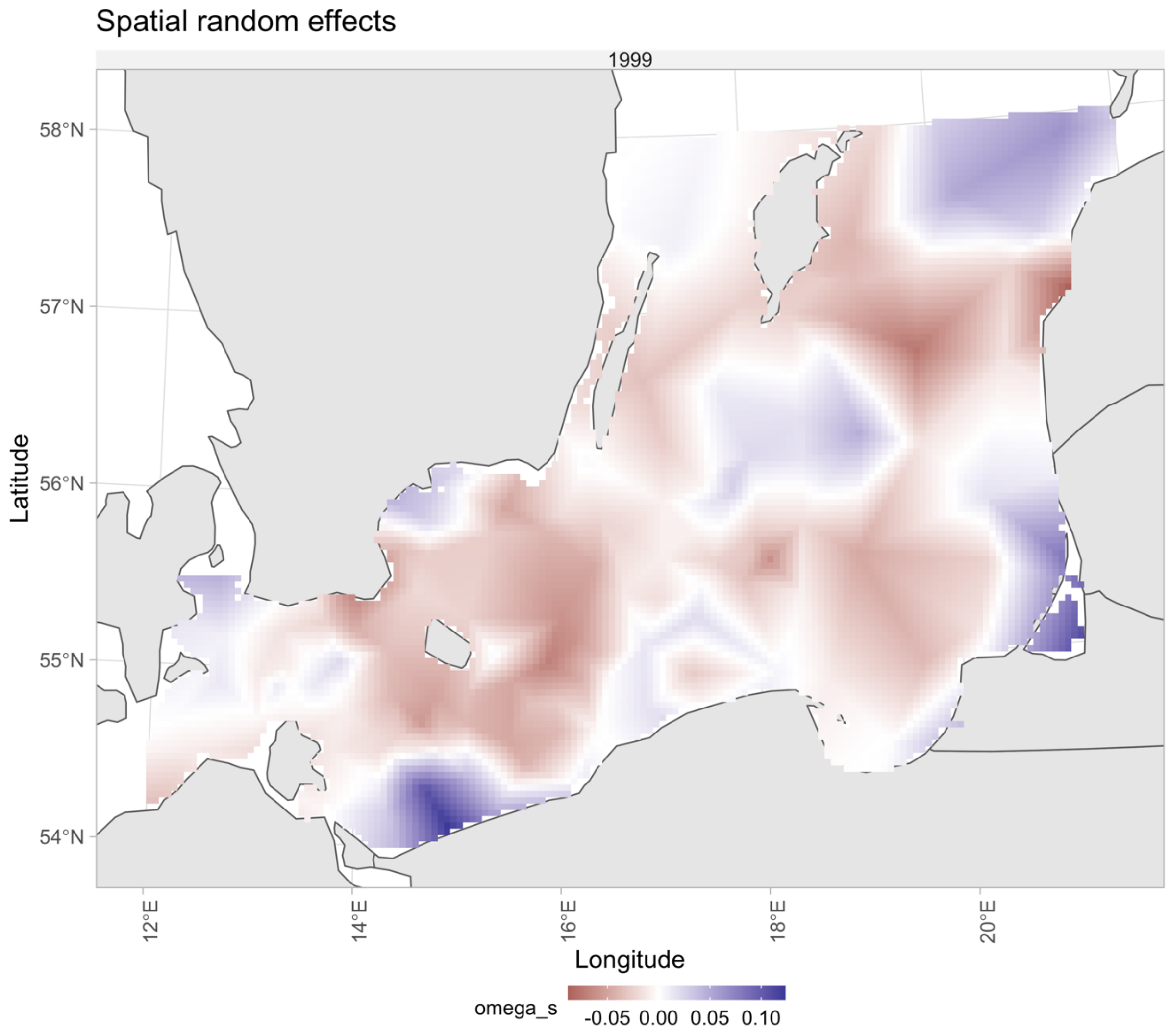


Fig. S8. Spatial random effects for the condition model.

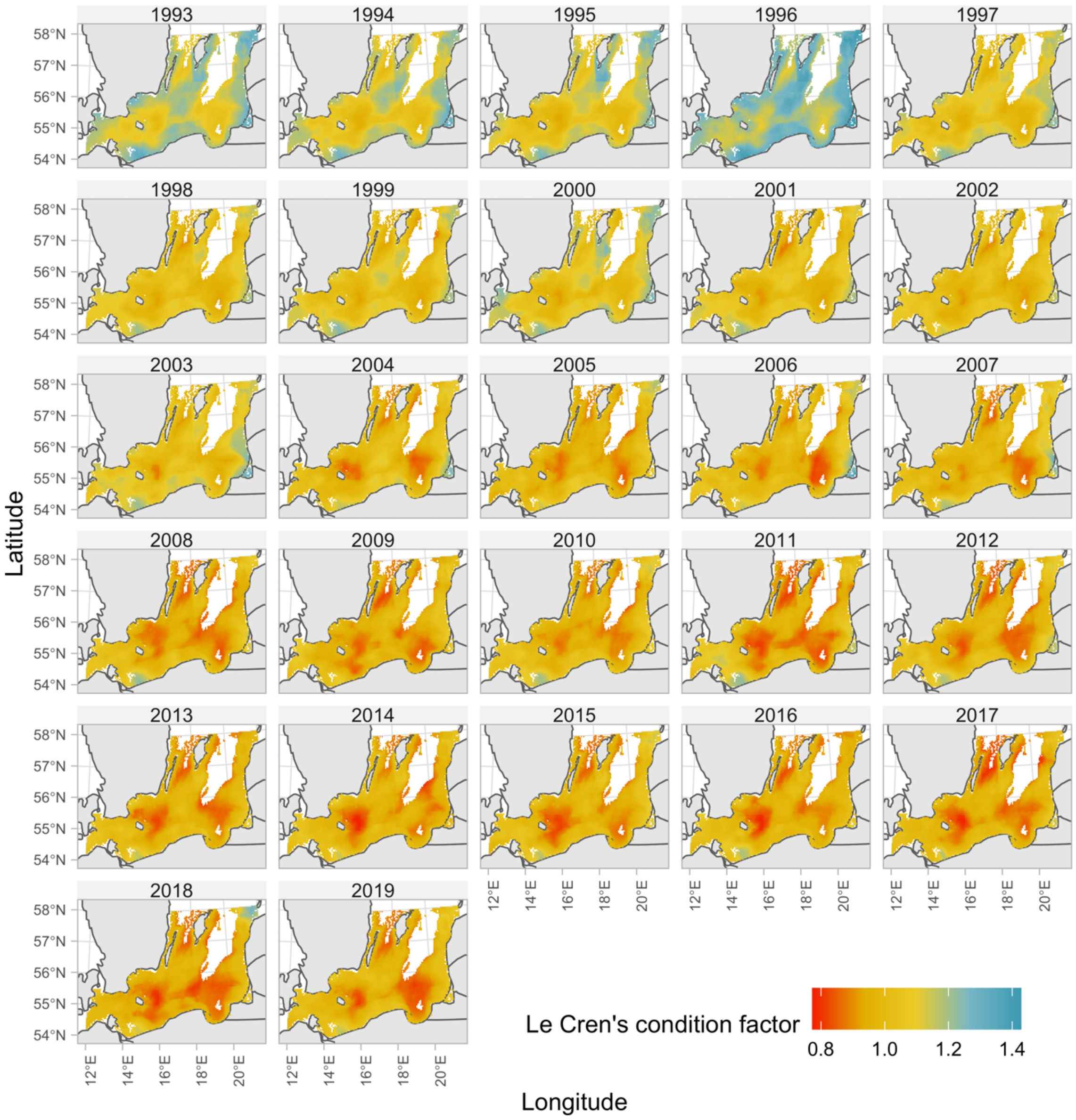


Fig. S9. Predicted condition factor with spatially varying covariates set to their true values (ICES rectangles with missing pelagic data were given the subdivision mean, see *SI Appendix*, Fig. S24).

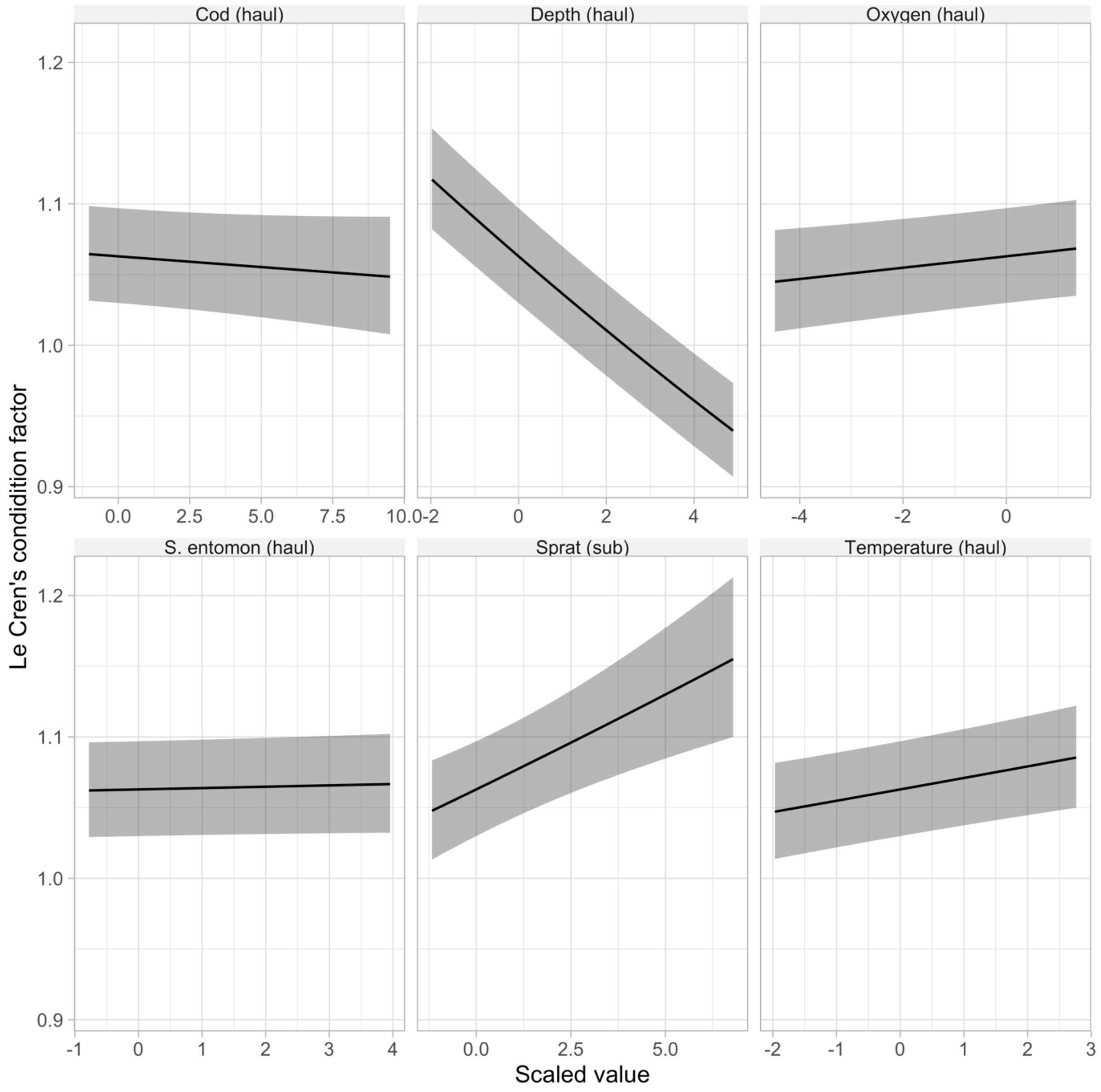


Fig. S10. Conditional effects from the condition model for selected covariates.

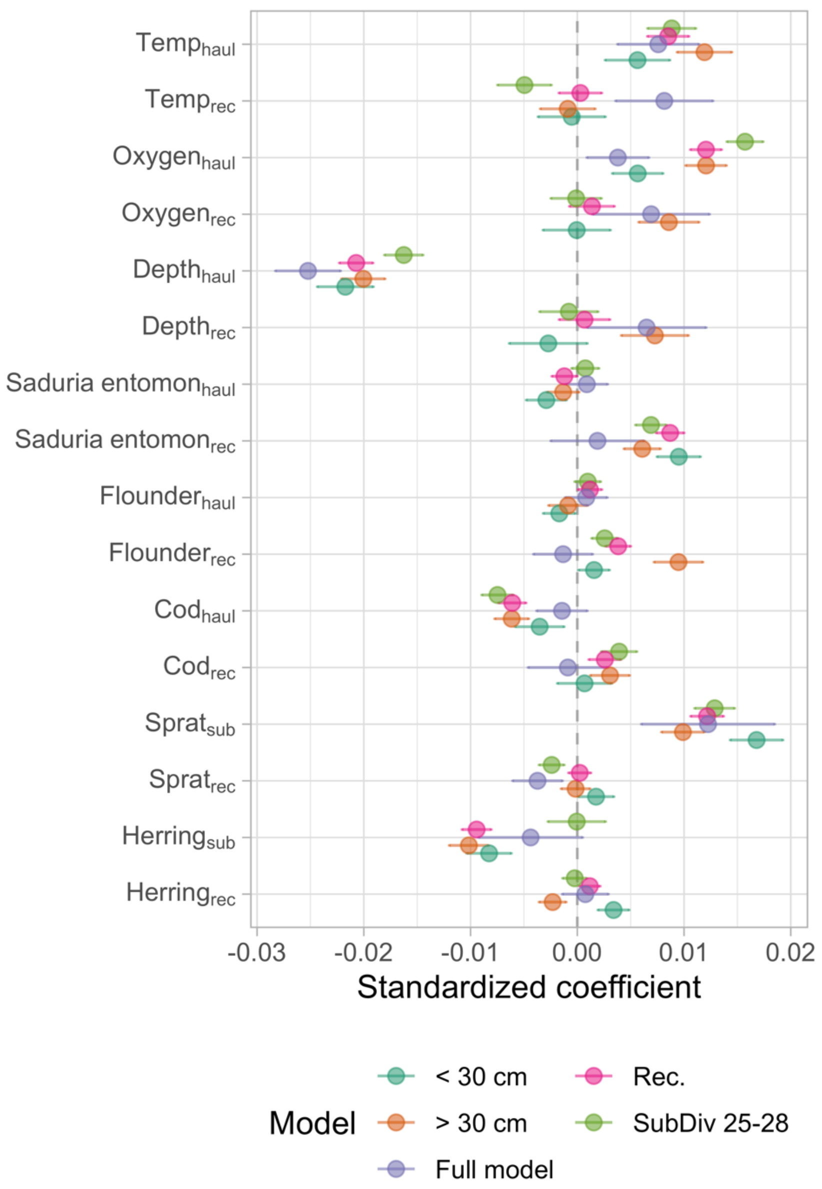


Fig. S11. Sensitivity analysis of the condition model. Each point corresponds to the covariate from a specific model, where the purple point is the model from the main text, teal is a model fitted only to cod below 30 cm, orange only above 30 cm (these models test if the coefficients are sensitive to the gradual ontogenetic diet shift cod exhibit). The green points stem from a model fitted to only subdivision 25-28, which corresponds to the core area of the eastern Baltic cod (subdivision 24 is a mixing zone between the distinct eastern and western Baltic cod). The pink points stem from a model where the rectangle-level medians of the covariates were calculated only using points on the grid with cod densities larger than the 5$^{th}$ percentile. Horizontal lines correspond to the 95% confidence interval).

Table S1. Comparison between models with oxygen as a linear effect (Eq. 6, main text) or as a linear breakpoint effect. The same model formulation was used as for the main model (Eqns. 1-4 in main text), but only covariates year (factor), oxygen (with or without threshold formula) and a linear effect for average oxygen in the ICES subdivision. The oxygen term is scaled (mean-centered and divided by the standard deviation). The breakpoint estimate of 1.07 corresponds to an oxygen concentration of 7.5 in units ml/L.

| **Model** | **Slope estimate (Standard error)** | **Breakpoint estimate** | **AIC** |
|---|---|---|---|
| Linear term | 0.0148 (0.0012) | NA | -144518.8 |
| Linear breakpoint term | 0.0088 (0.0017) | 1.0700 (0.3740) | -144450.5 |

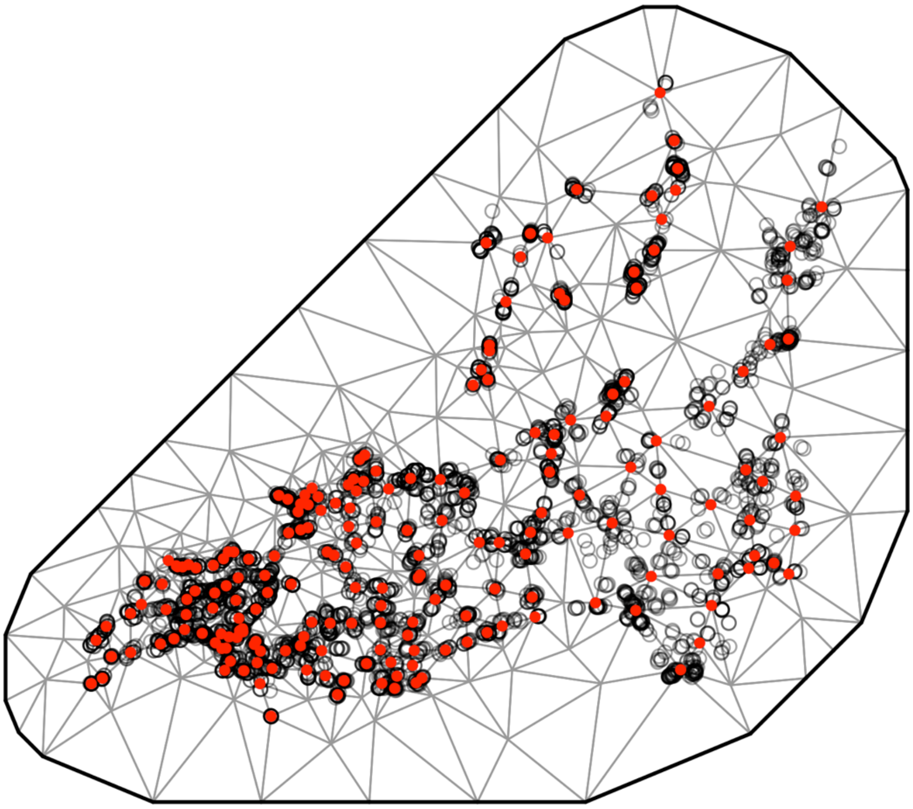

Fig. S12. SPDE mesh for the cod density model (200 knots).

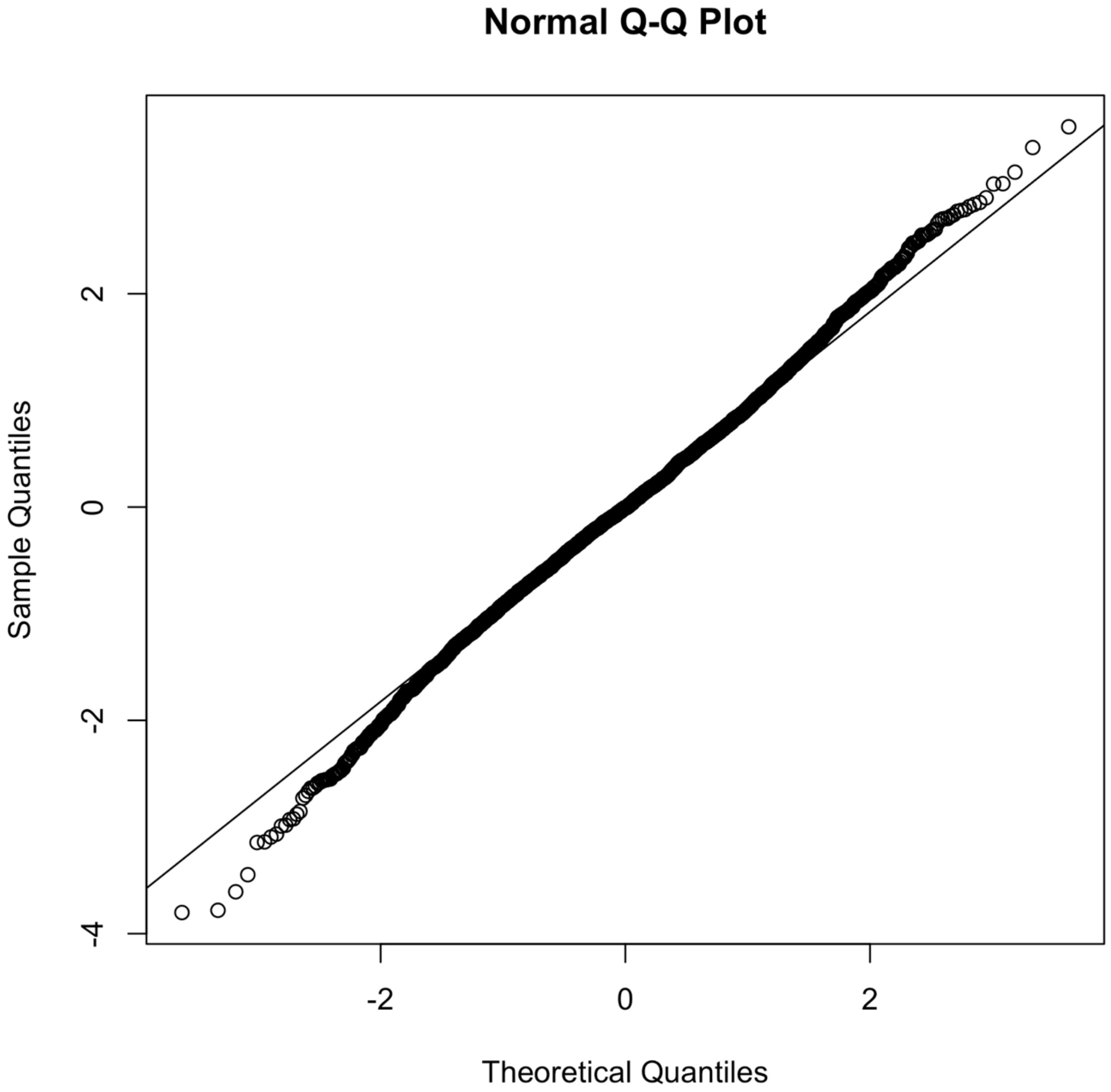


Fig. S13. QQ plot of cod density model based on randomized quantile residuals (fixed effects held at their maximum likelihood estimate and the random effects are sampled with MCMC via ‘tmbstan’ [1] and ‘Stan’ [2], in line with [3,4].

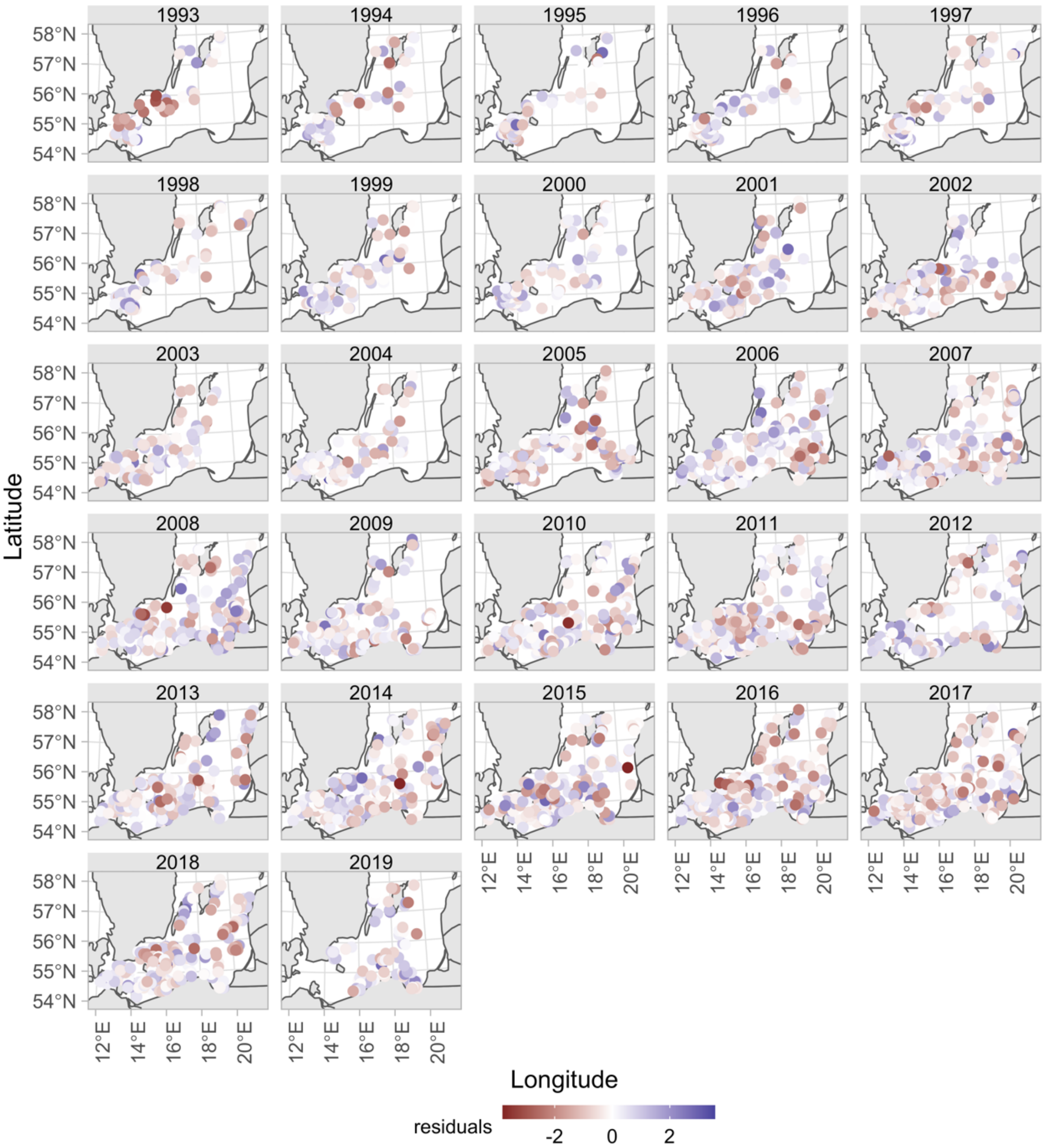


Fig. S14. Cod density model residuals plotted in space

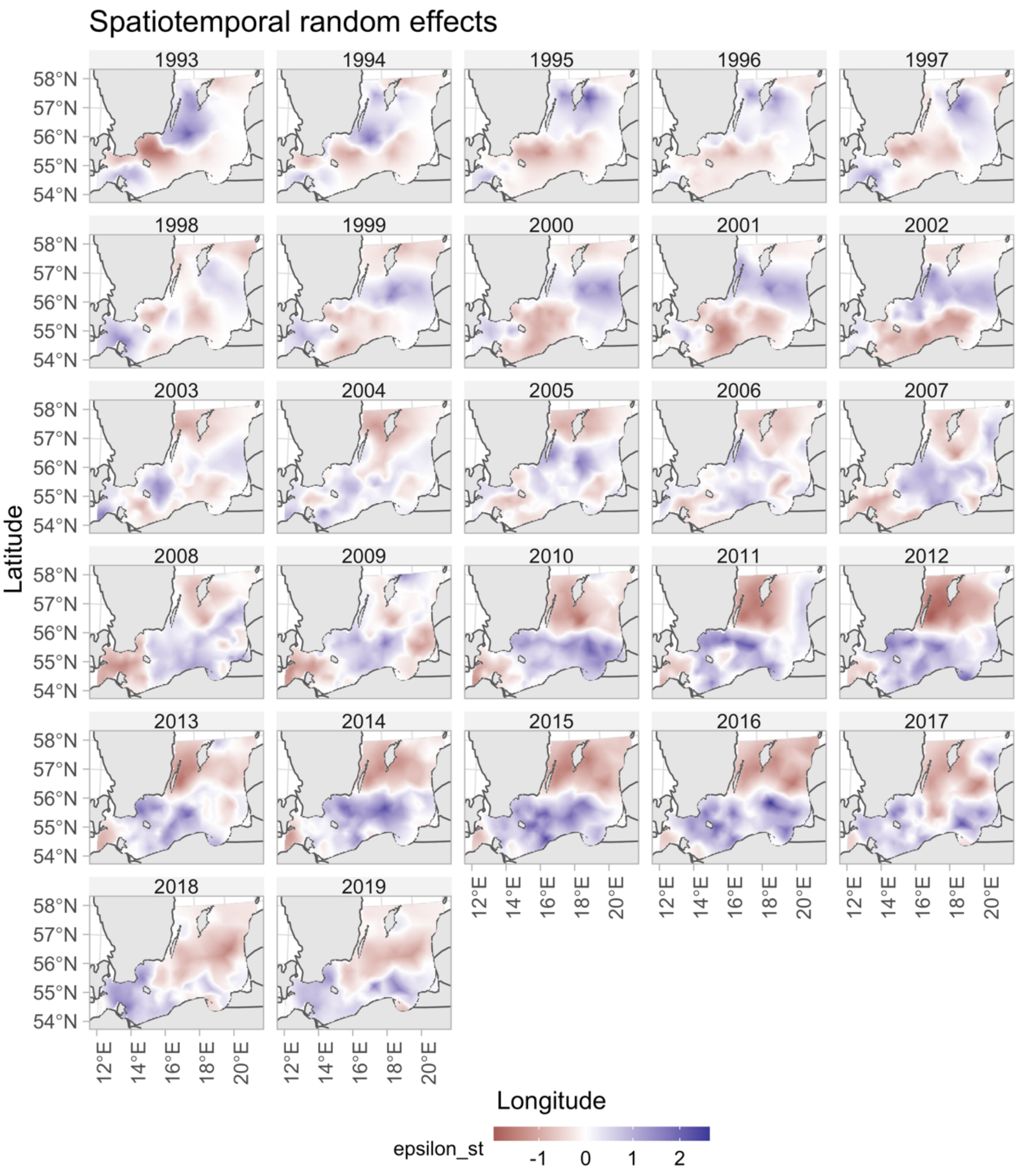


Fig. S15. Spatiotemporal random effects for the cod density model

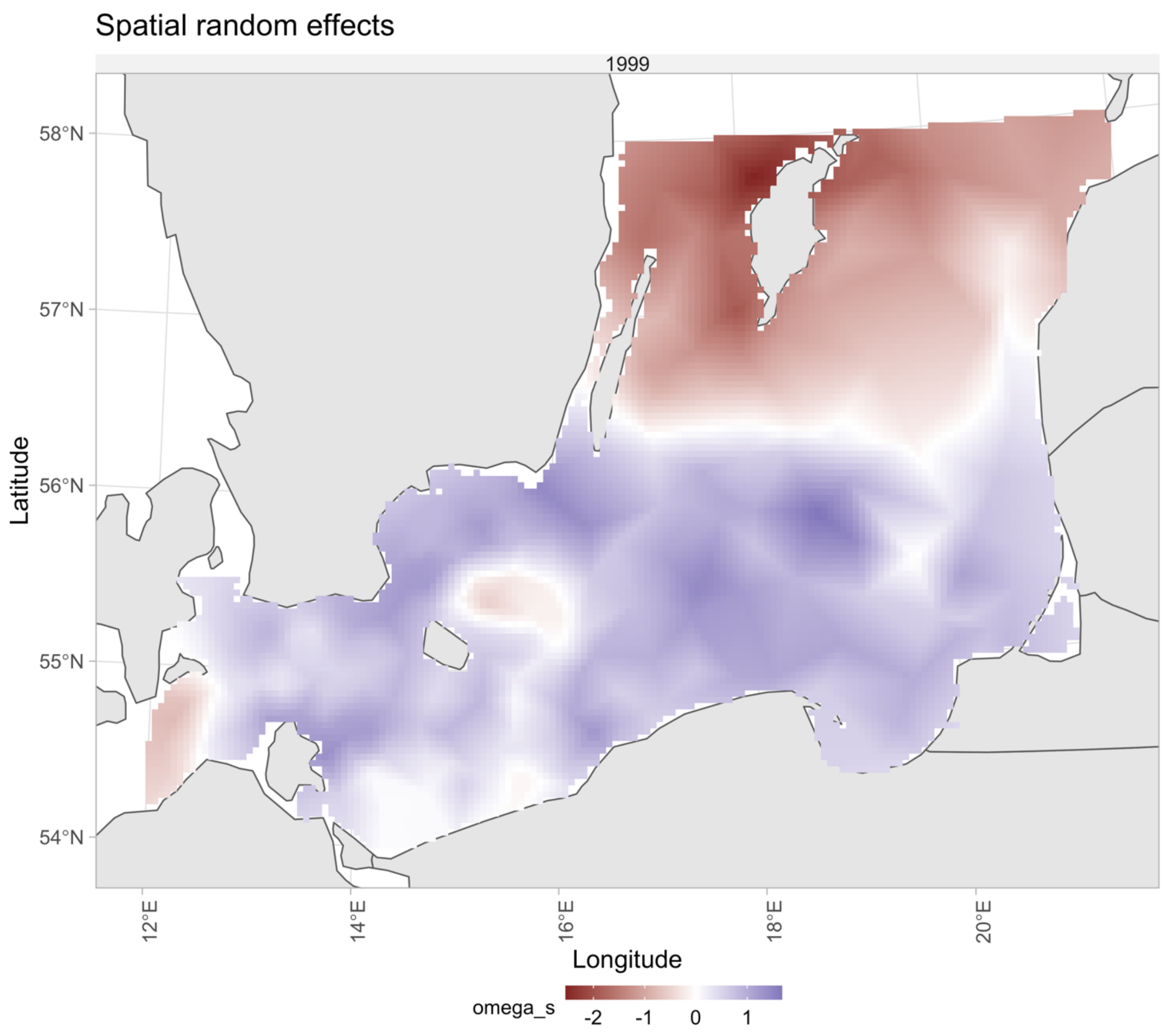


Fig. S16. Spatial random effects for the cod density model.

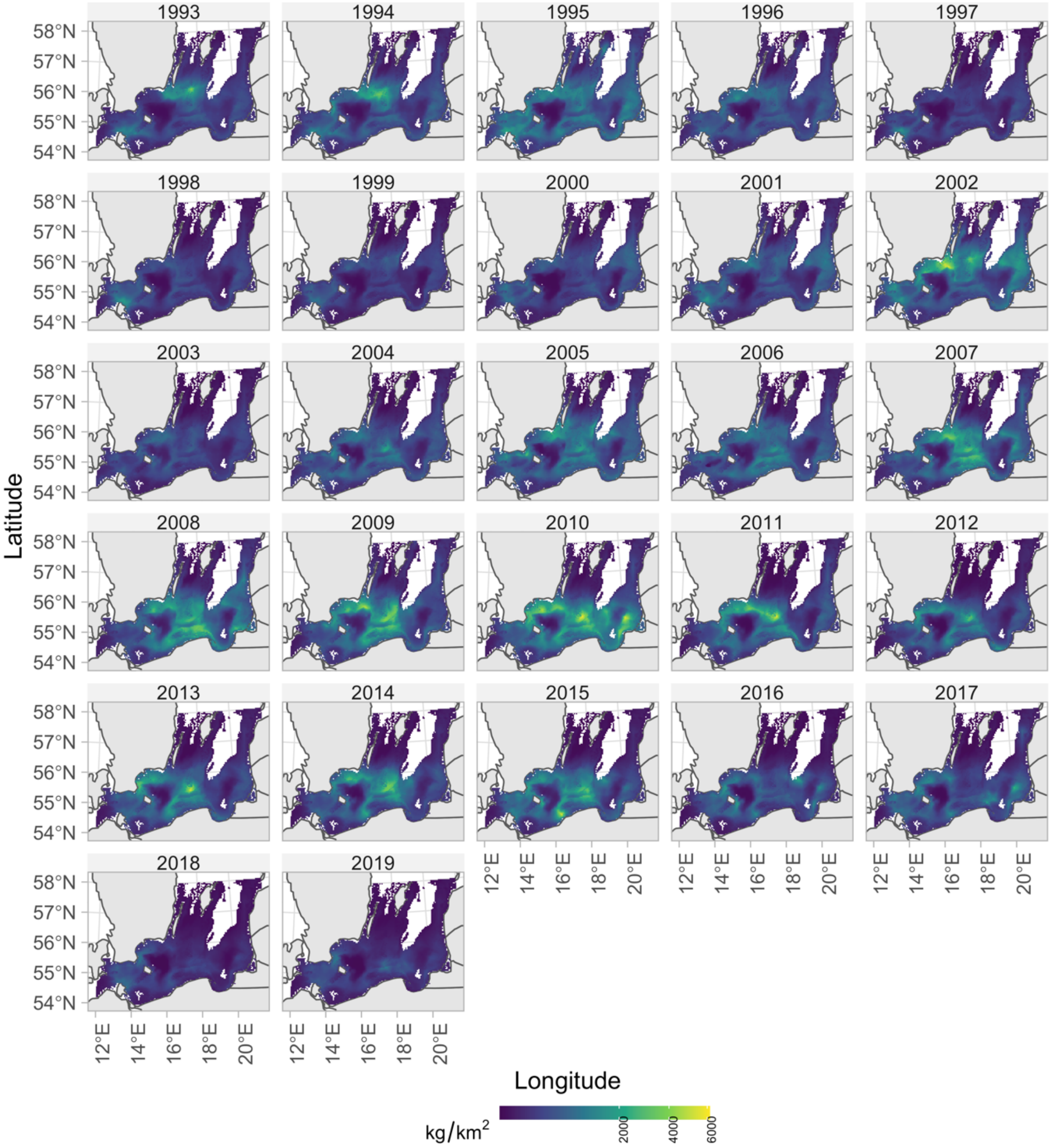


Fig. S17. Predicted cod density in space and time with covariates depth, oxygen and temperature.

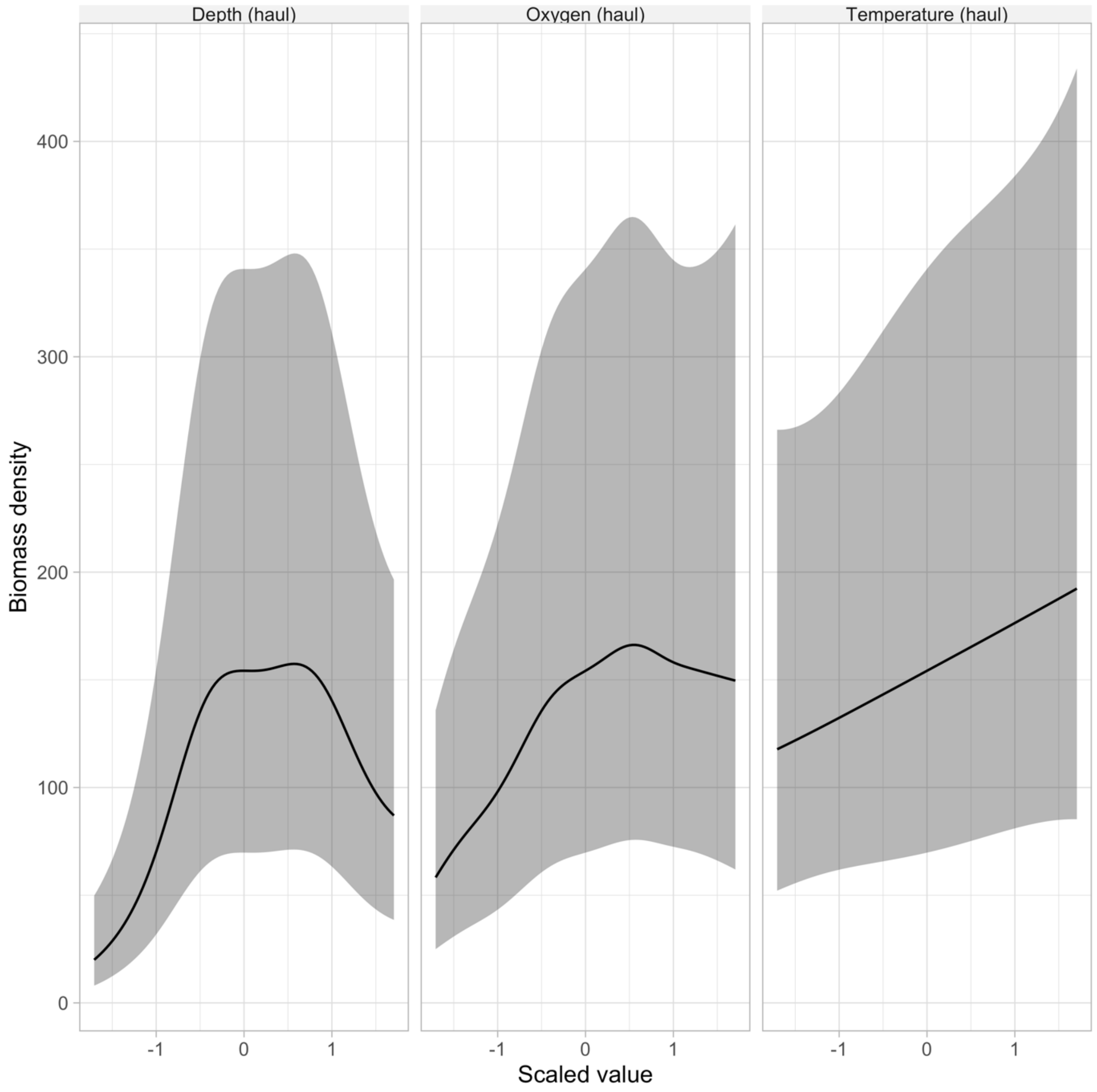


Fig. S18. Conditional effects from the cod density model

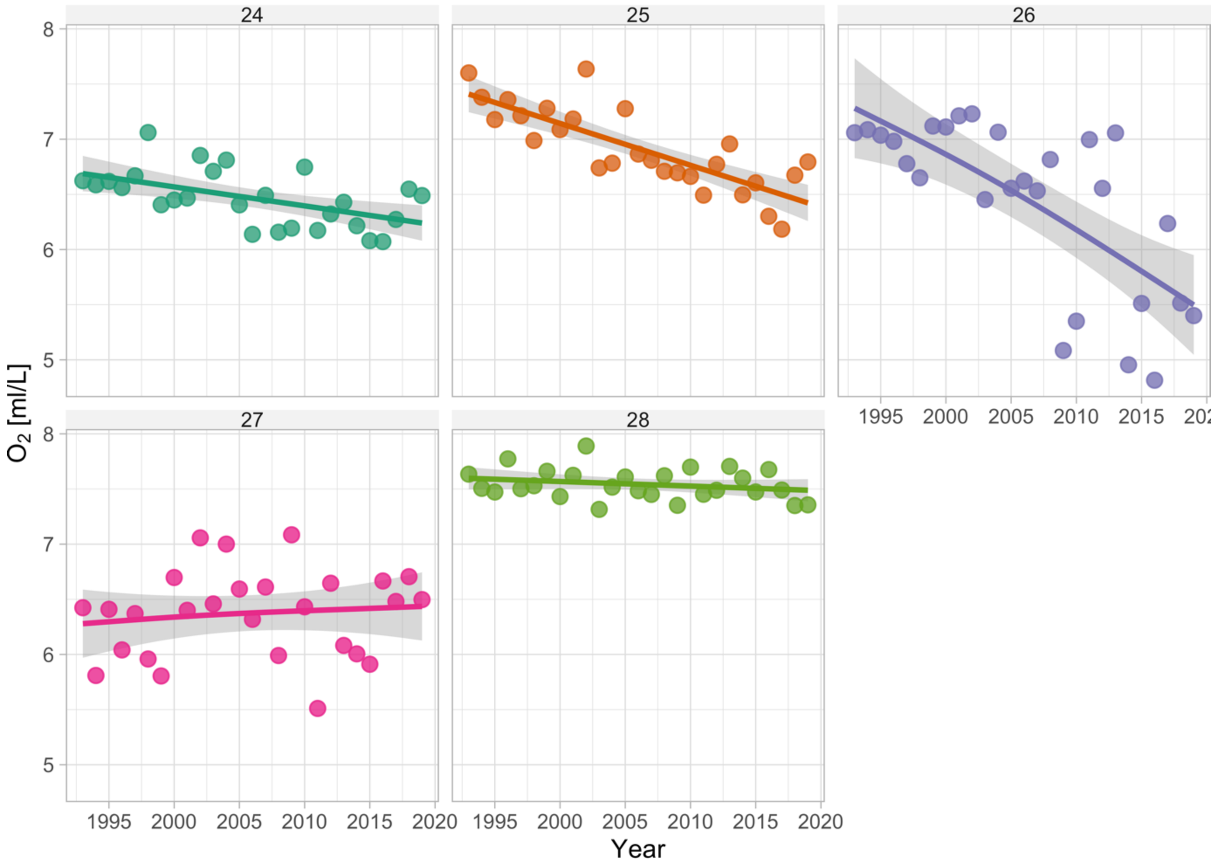


Fig. S19. Density-weighted sea bottom oxygen by sub-division. Lines depict GAM fits ($k$=4) fits and the shaded area depicts the 95% confidence intervals.

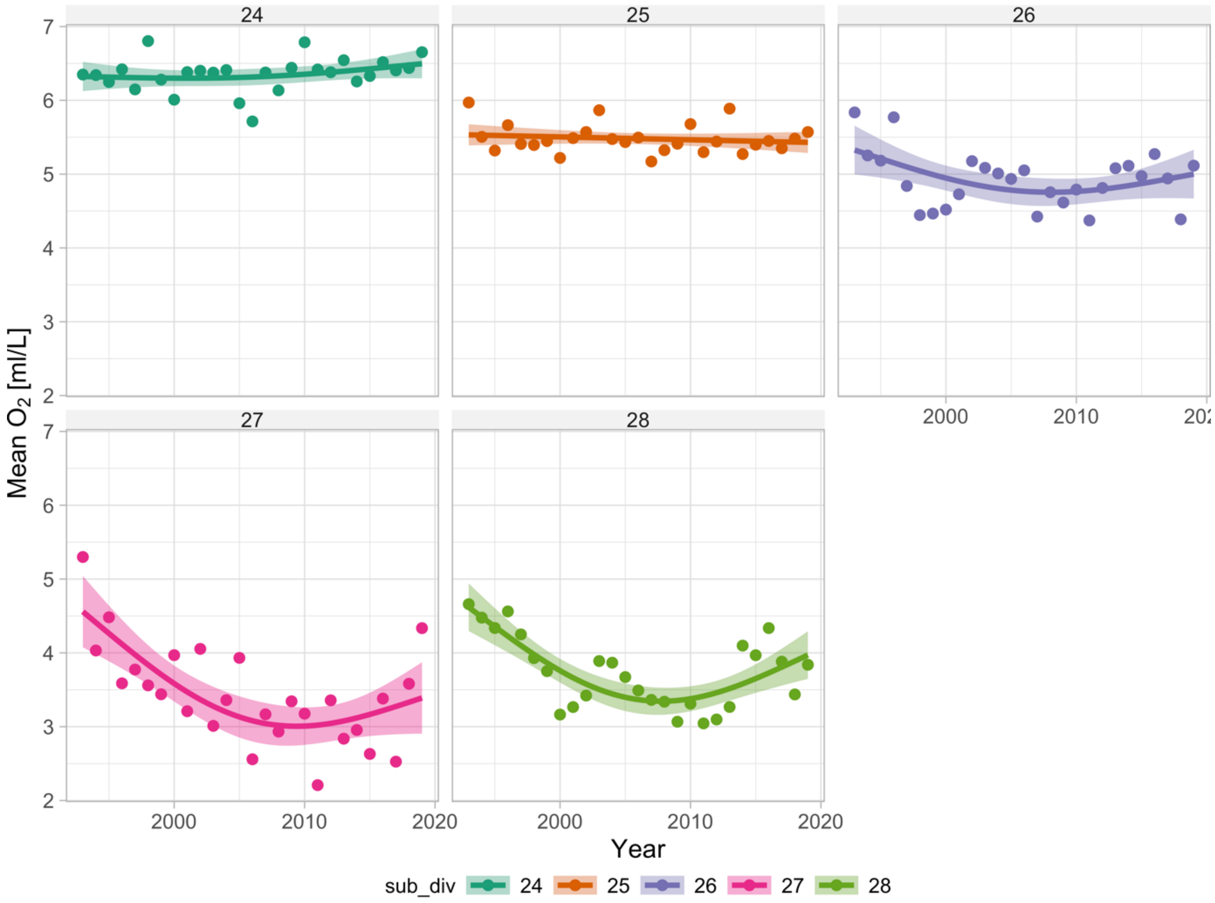


Fig. S20. Sea bottom oxygen concentration in the environment, by sub-division. Lines depict GAM fits (*k*=4) fits and the shaded area depicts the 95% confidence intervals.

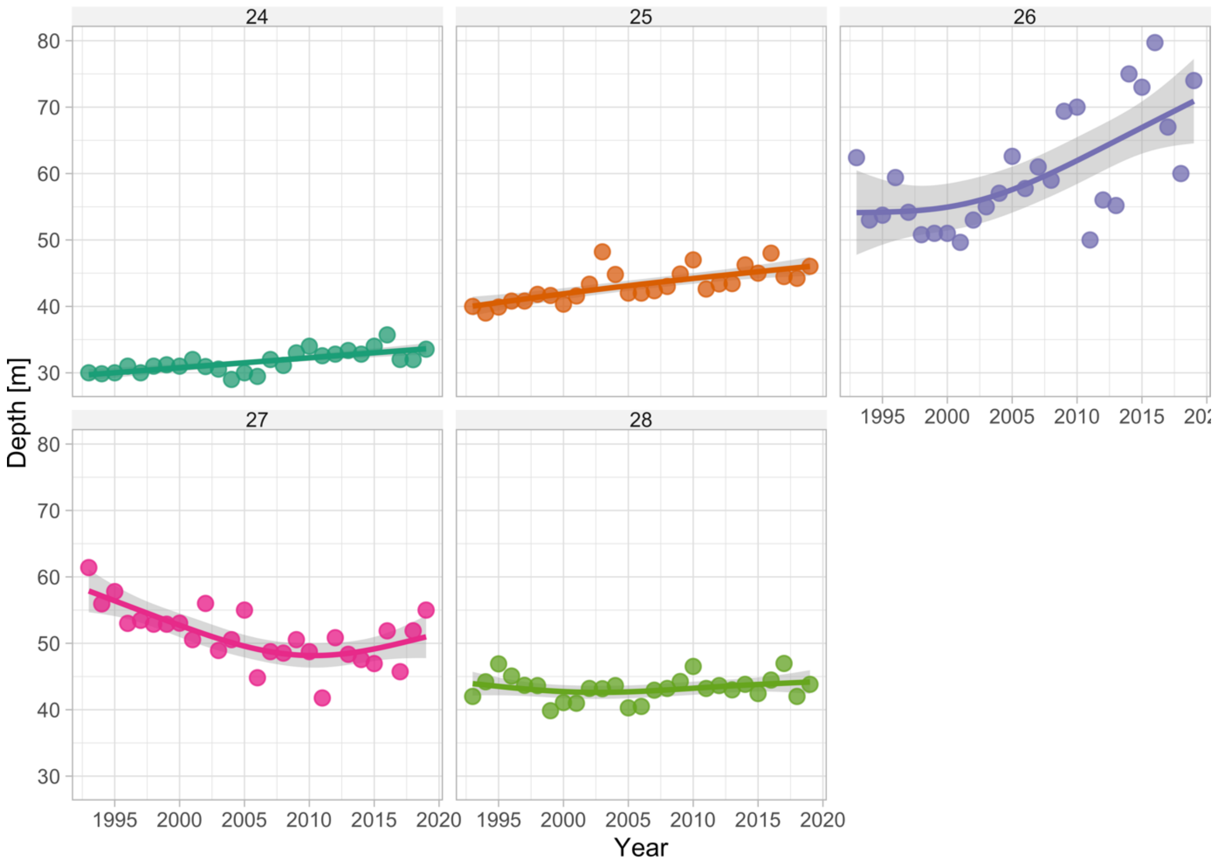


Fig. S21. Density-weighted depth by sub-division. Lines depict GAM fits ($k$=4) fits and the shaded area depicts the 95% confidence intervals.

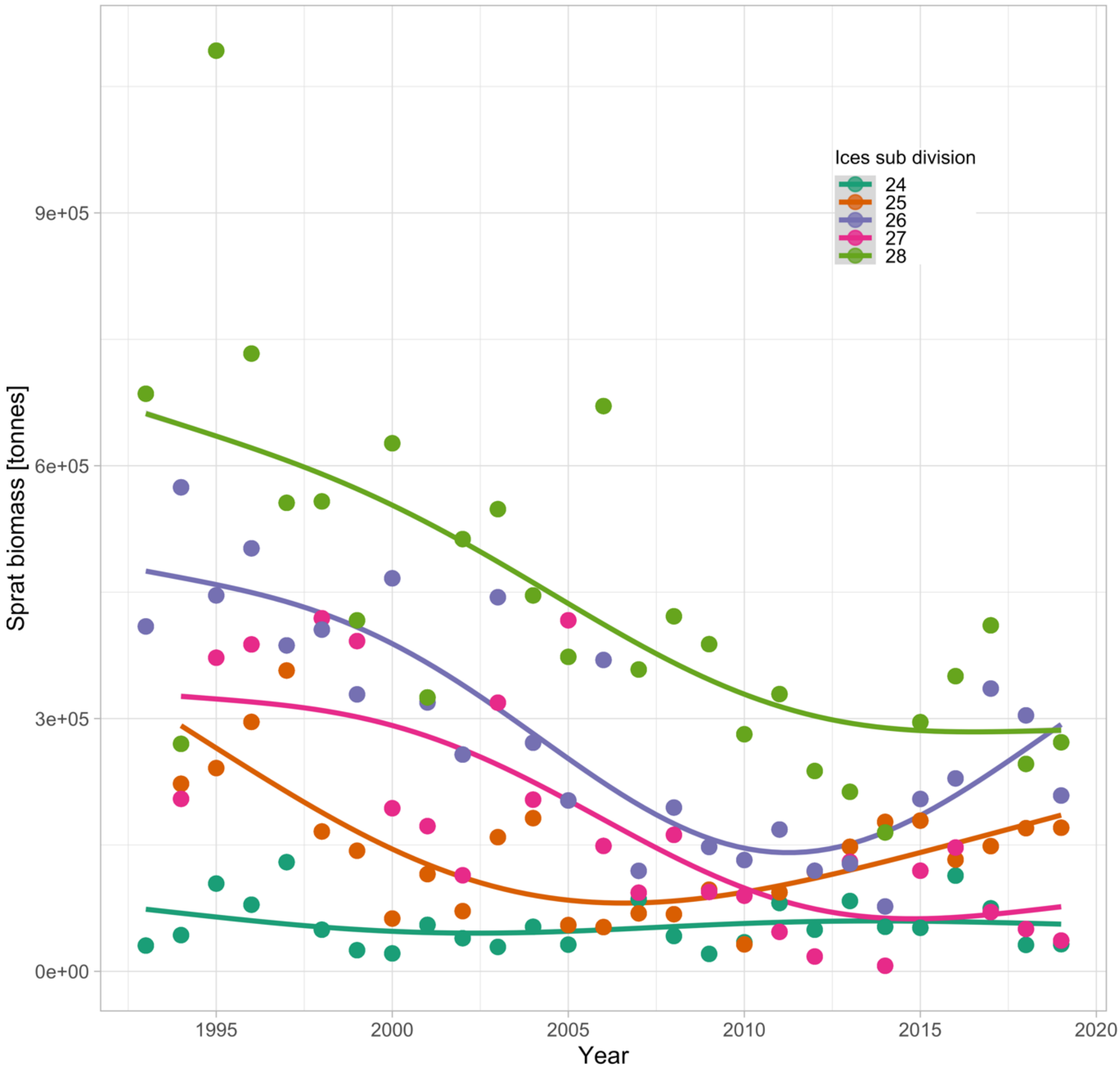


Fig. S22. Biomass of sprat over time by sub-division. Lines depict GAM fits (*k*=4) fits.

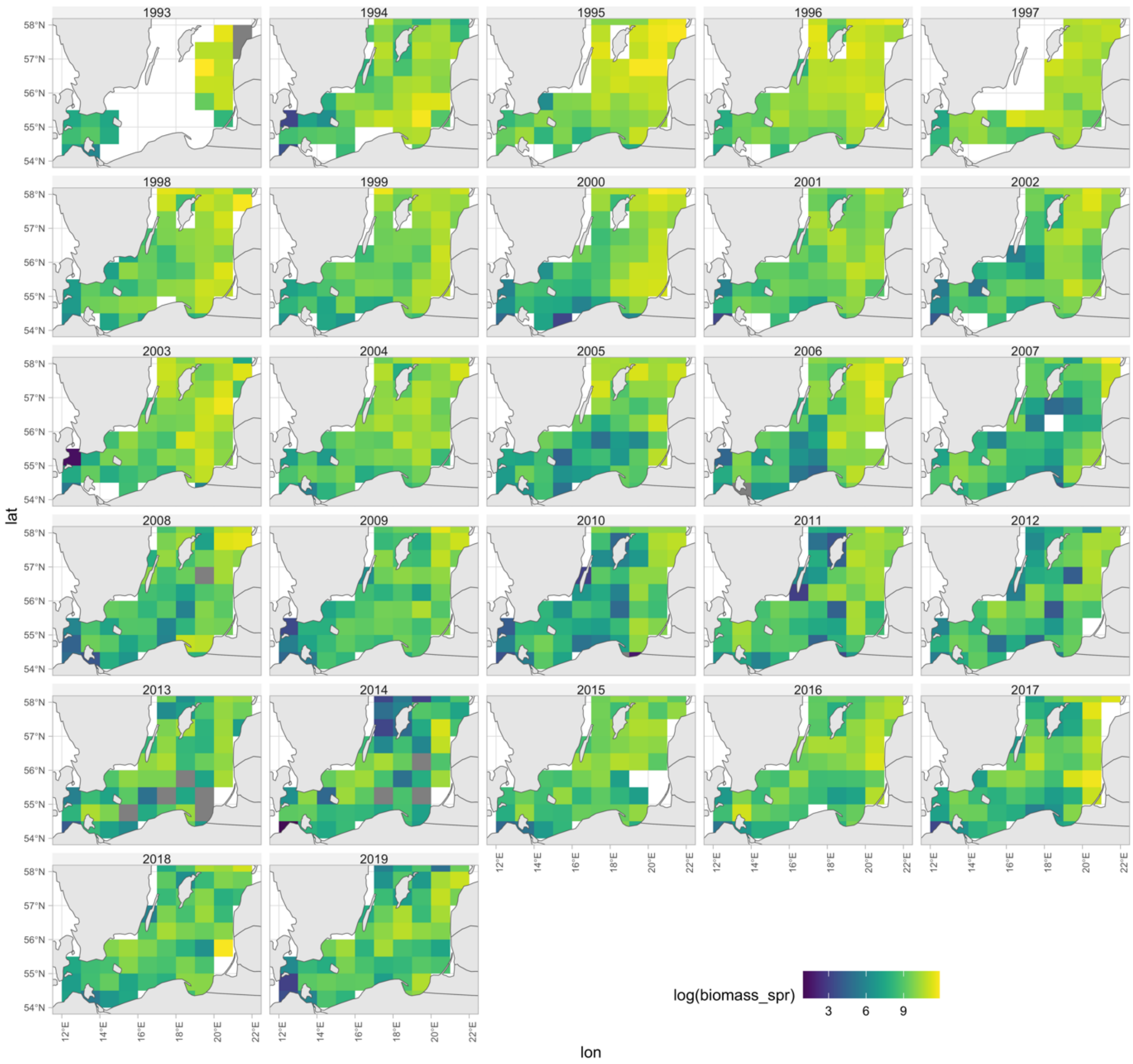


Fig. S23. Rectangle-level log biomass of sprat indicating rectangles with missing values.

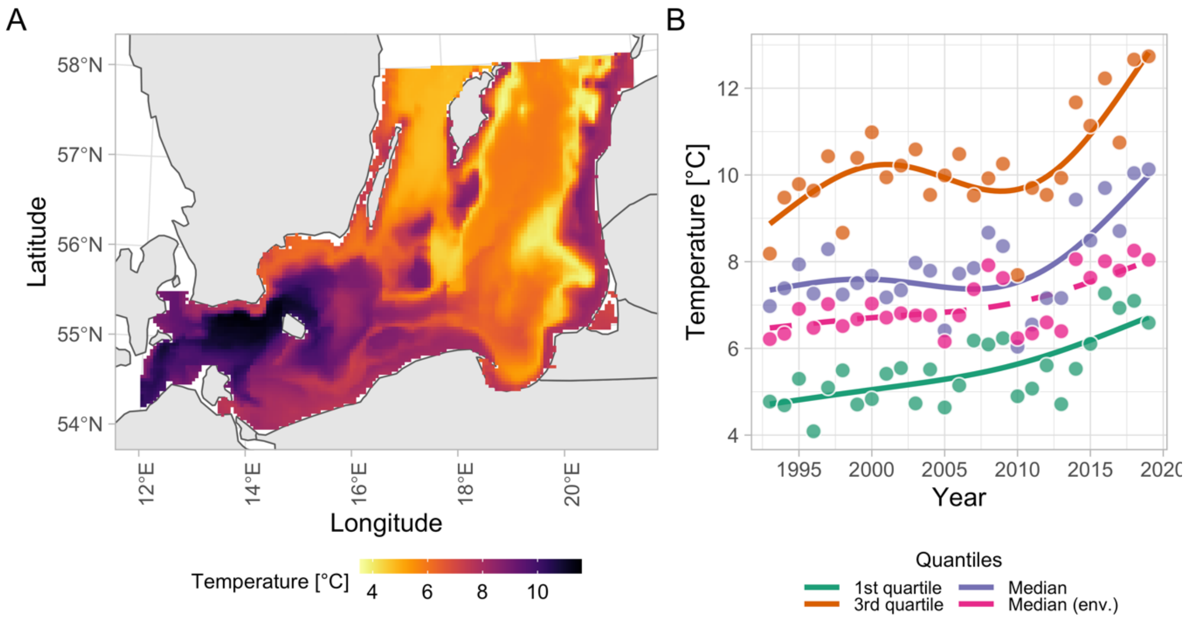


Fig. S24. Sea bottom temperature (exemplified using year 2006) in the study area. Panel (B) temperature weighted by predicted cod density. Colors indicate quantiles (1st quartile, median and 3rd quartile). Lines depict GAM fits (*k*=4).

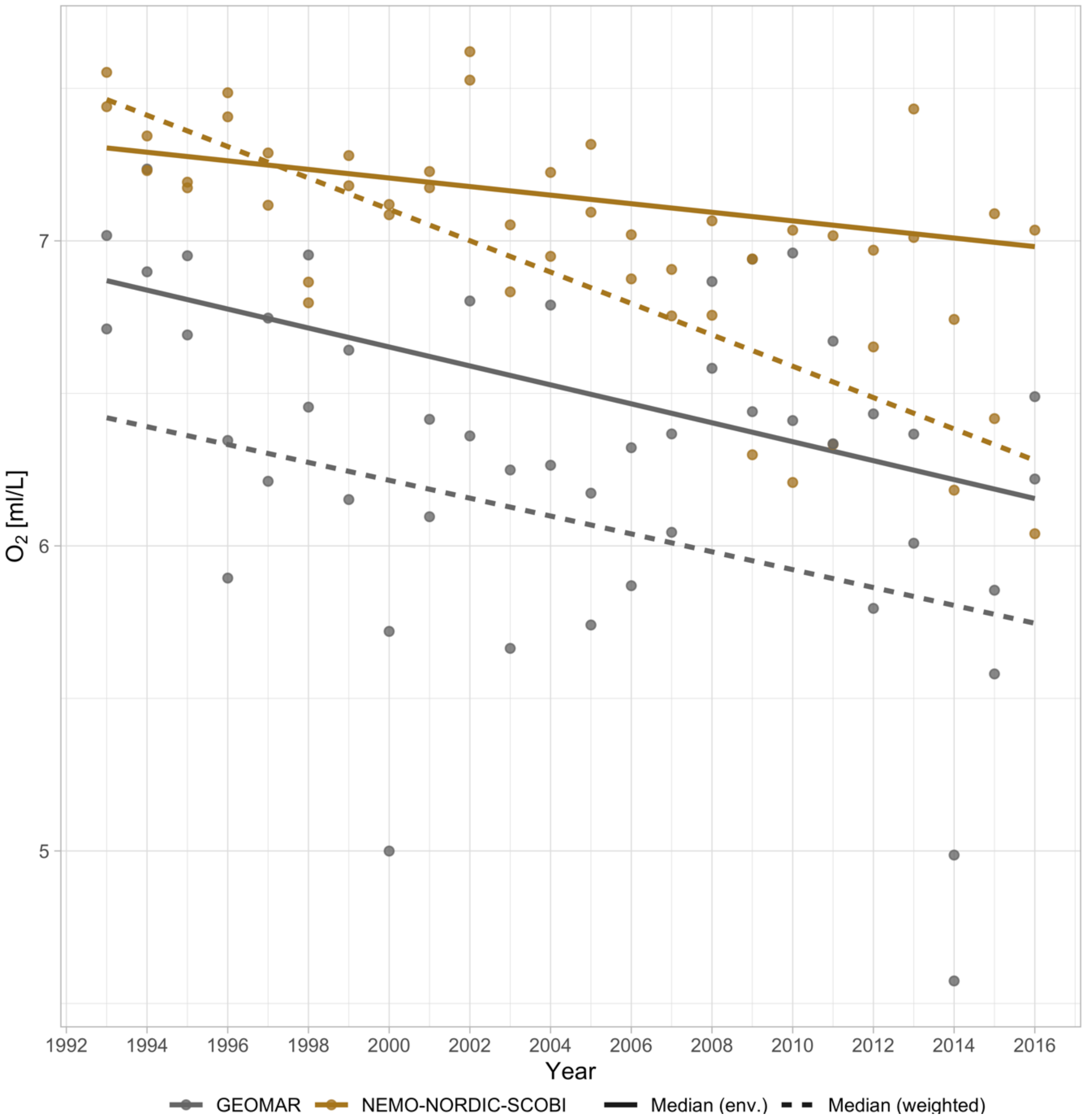


Fig. S25. Comparison of oxygen concentrations between the "GEOMAR" model used in Lehmann et al., [5,6] (gray lines and points) and the [7–9] NEMO-NORDIC-SCOBI models (brown lines and points) in ICES subdivision 25 between 1993–2016. Solid lines depict the median in the environment in the interquartile depth range (29-61 m), and dashed lines depict the median oxygen concentration weighted by the predicted biomass density of cod.
